# Mutations in the B.1.1.7 SARS-CoV-2 spike protein reduce receptor-binding affinity and induce a flexible link to the fusion peptide

**DOI:** 10.1101/2021.04.06.438584

**Authors:** Eileen Socher, Marcus Conrad, Lukas Heger, Friedrich Paulsen, Heinrich Sticht, Friederike Zunke, Philipp Arnold

**Affiliations:** Institute of Anatomy, Functional and Clinical Anatomy, Friedrich-Alexander-University Erlangen-Nürnberg (FAU), Erlangen, Germany; Institute for Clinical and Molecular Virology, Friedrich-Alexander-University Erlangen-Nürnberg (FAU), University Hospital Erlangen, Erlangen, Germany; Division of Bioinformatics, Institute of Biochemistry, Friedrich-Alexander-University Erlangen-Nürnberg (FAU), Erlangen, Germany; Laboratory of Dendritic Cell Biology, Department of Dermatology, Friedrich-Alexander-University Erlangen-Nürnberg (FAU), University Hospital Erlangen, Erlangen, Germany; Sechenov University, Department of Operative Surgery and Topographic Anatomy, Moscow, Russia; Erlangen National High Performance Computing Center (NHR@FAU), Friedrich-Alexander-University Erlangen-Nürnberg (FAU), Erlangen, Germany; Department of Molecular Neurology, University Hospital Erlangen, Friedrich-Alexander-University Erlangen-Nürnberg (FAU), Erlangen, Germany

**Keywords:** COVID-19, SARS-CoV-2, B.1.1.7, molecular dynamics simulation, ACE2, receptor binding

## Abstract

The B.1.1.7 variant of the SARS-CoV-2 virus shows enhanced infectiousness over the wild type virus, leading to increasing patient numbers in affected areas. A number of single amino acid exchanges and deletions within the trimeric viral spike protein characterize this new SARS-CoV-2 variant. Crucial for viral entry into the host cell is the interaction of the spike protein with the cell surface receptor angiotensin-converting enzyme 2 (ACE2) as well as integration of the viral fusion peptide into the host membrane. Respective amino acid exchanges within the SARS-CoV-2 variant B.1.1.7 affect inter-monomeric contact sites within the spike protein (A570D and D614G) as well as the ACE2-receptor interface region (N501Y), which comprises the receptor-binding domain (RBD) of the viral spike protein. However, the molecular consequences of mutations within B.1.1.7 on spike protein dynamics and stability, the fusion peptide, and ACE2 binding are largely unknown. Here, molecular dynamics simulations comparing SARS-CoV-2 wild type with the B.1.1.7 variant revealed inter-trimeric contact rearrangements, altering the structural flexibility within the spike protein trimer. In addition to reduced flexibility in the N-terminal domain of the spike protein, we found increased flexibility in direct spatial proximity of the fusion peptide. This increase in flexibility is due to salt bridge rearrangements induced by the D614G mutation in B.1.1.7 found in pre- and post-cleavage state at the S2’ site. Our results also imply a reduced binding affinity for B.1.1.7 with ACE2, as the N501Y mutation restructures the RBD-ACE2 interface, significantly decreasing the linear interaction energy between the RBD and ACE2.

Our results demonstrate how mutations found within B.1.1.7 enlarge the flexibility around the fusion peptide and change the RBD-ACE2 interface, which, in combination, might explain the higher infectivity of B.1.1.7. We anticipate our findings to be starting points for in depth biochemical and cell biological analyses of B.1.1.7, but also other highly contagious SARS-CoV-2 variants, as many of them likewise exhibit a combination of the D614G and N501Y mutation.

## Introduction

The outbreak of the severe acute respiratory syndrome (SARS) caused by the SARS-like coronavirus SARS-CoV-2 has emerged to a global pandemic with daily increasing numbers of infections and deaths exceeding 2.5 million world-wide^1^ On top of the rapid, global spread of SARS-CoV-2, new and more contagious virus variants comprise an additional threat^2–5^. The novel SARS-CoV-2 variant B.1.1.7 first emerged in southeast England in November 2020 and is estimated to be 56% more transmissible^6^. This variant is characterized by a number of amino acid deletions and exchanges, with most of the protein-coding mutations found within the surface-anchored spike (S) protein of the virus: del69–70HV, del144Y, N501Y, A570D, D614G, P681H, T761I, S982A, D1118H^7^ (sFig1 a). The homotrimeric S protein facilitates viral entry into host cells by interaction of its receptor-binding domain (RBD) with the cell surface receptor angiotensin-converting enzyme 2 (ACE2)^7–9^. The S protein consist of a S1 subunit, harboring the receptor-binding domain, and a S2 subunit, that is essential for viral entry into the host cell after dissociation from S1 (Fig1 a, sFig1b). Separation of S1 and S2 subunit is mediated by host cell surface proteases at the S1/S2 and S2’ cleavage sites of the S protein^10, 11^. Previous results suggest that dramatic structural changes of the S2 unit lead to the exposure of a fusion peptide that facilitates cell membrane fusion and virus entry^12^. For S protein priming, human serin protease TMPRSS2 cleavage within a multi-basic (furin) cleavage site between AAs685/686 (S1/S2 cleavage site) or AAs815/816 (S2’ cleavage site) has been proposed^9, 13^ and a recent study even suggests blocking SARS-CoV-2 infection by utilizing a TMPRSS2 inhibitor^9^.

**Figure 1:**
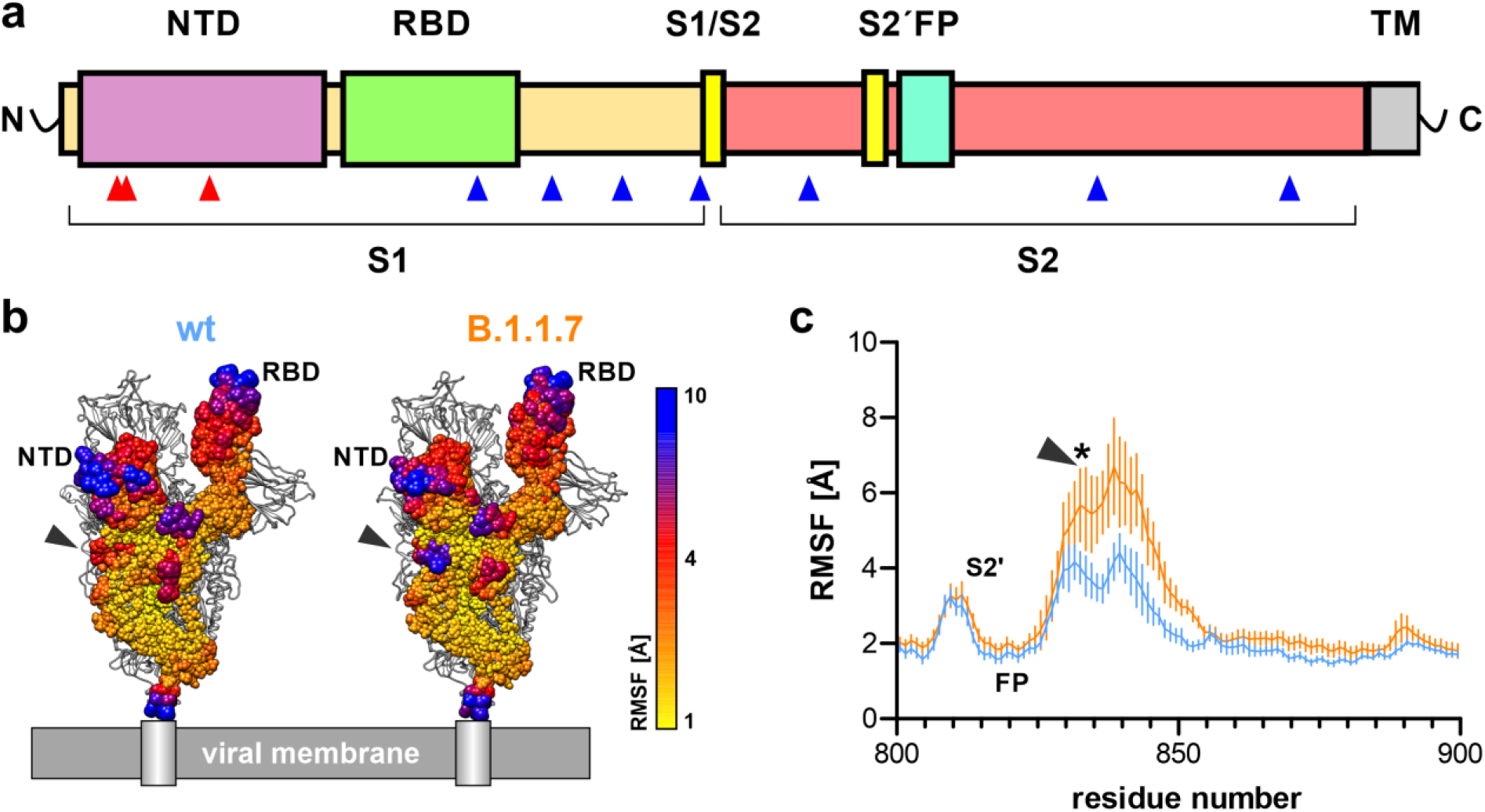
Structural flexibility in the B.1.1.7 SARS-CoV-2 S protein. a) Schematic primary structure of the SARS-CoV-2 S protein indicating the site of amino acid deletions (red arrowheads) or single amino acid exchanges (blue arrowheads; NTD, N-terminal domain; RBD, receptor-binding domain; S1/S2, furin cleavage site at positions 685/686; S2’, furin/TMPRSS2 cleavage site at positions 815/816; FP, fusion peptide; TM, transmembrane domain and C-terminal end). b) S protein trimer as it would reside on the cell surface with one subunit colored for structural flexibility as calculated during simulation (root-mean-square fluctuations (RMSF), n = 6). NTD denotes the N-terminal domain, RBD the receptor-binding domain and the grey arrowhead the loop region between amino acids 835-843. c) Line plot of RMSF values for amino acid residues 800-900 reveals increased flexibility for residues 835 and 843 in B.1.1.7 (orange) when compared to wt (blue). The arrowhead denotes the same region as in b) and the asterisk indicates statistical differences for these amino acids (n = 6; two-way ANOVA; statistical significance assumed for *p<0.05; full statistic can be found in Tab. 2).

Cellular uptake as well as protein structure of the S protein of the SARS-CoV-2 virus have been studied extensively. Hence, cryo-electron microscopy and crystallographic analyses of the S protein trimer^14, 15^, ACE-2-bound S protein^16^ as well as ACE-2-bound receptor-binding domain^17^ have shed light onto the structural mechanisms of viral entry. Until now this structural information is missing for globally emerging SARS-CoV-2 variants, exhibiting several mutations within the S protein and higher infectivity. We only have a vague understanding of the molecular reasons leading to enhanced contagion and cellular uptake of SARS-CoV-2 variants like the B.1.1.7 (British), but also the B.1.351 (South African) and P.1 (Brazilian/Japanese) variants. Interestingly, these three Variants of Concern (VOC) have the N501Y and D614G mutation in common^2, 18^

In the present study, we focus on the B.1.1.7 variant and utilize molecular dynamics (MD) simulations of S protein trimer to assess its dynamic behavior in terms of conformational stability as well as interaction of isolated viral RBD in complex with human ACE2 to calculate linear interaction energies for wild type and mutated (N501Y) RBD-ACE2 complexes. Our data suggest increased flexibility around the fusion peptide induced by the D614G mutation and reduced binding affinity between RBD and ACE2 due to conformational reorganization of the RBD-ACE2 interface mediated by N501Y. Upon cleavage at the S2’ cleavage site (Arg815/Ser816), a conformational switch alters partial salt bridge formation between arginine 815 and negatively charged aspartate and glutamate residues in the vicinity to keep the new C-terminus in place via non-covalent bindings.

## Results

### Flexibility of the spike trimer

To compare flexibility between the SARS-CoV-2 wild type (wt) and B.1.1.7 variant, root-mean-square fluctuation (RMSF) values were calculated for the backbone atoms of all individual residues over 200 ns simulation time, averaged for all six calculated monomers (two runs with three subunits per trimer) and visualized as spheres of different diameter and color according to their average RMSF values (Fig1 b, sFig2 a,b). Highest RMSF values were calculated for the RBD with flexibility of up to 10 Å especially at the interface positions where interaction with ACE2 would occur upon receptor binding (sFig2 a,b,c). In the N-terminal domain (NTD) deletions of amino acids 69, 70 and 144 induce a reduced flexibility for amino acid residues arginine 78, leucine 249 and threonine 250 in the B.1.1.7 variant (sFig2 d, Tab. 1). These residues are all located at the surface of the spike protein and are most exposed to interact with other spike trimers at the viral surface (sFig2 b,e). The loop region C-terminal of the S2’ cleavage site and the fusion peptide showed increased flexibility in the B.1.1.7 variant when compared to wt (Fig1 c, sFig2 c). Flexibility increased from a maximum of 4 Å in wt to a maximum of about 7 Å in B.1.1.7. Residues 835-843 showed markedly increased RMSF values (Tab. 2). To understand this change in flexibility we analyzed the salt bridges formed by charged residues of this amino acid stretch.

**Tab.1:** Statistical analysis of rmsf plots from wt and B.1.1.7 spike protein residues 26-300 (n =6; two-way ANOVA; significance assumed for *p<0.05).

| Compare each cell mean with the other cell mean in that row |  |  |  |  |  |
| --- | --- | --- | --- | --- | --- |
| Number of families | 1 |  |  |  |  |
| Number of comparisons per family | 268 |  |  |  |  |
| Alpha | 0,05 |  |  |  |  |
| Sidak's multiple comparisons test | Mean Diff, | 95,00% CI of diff, | Significant? | Summary | Adjusted P Value |
| residue number |  |  |  |  |  |
| 26 | -0,8329 | -2,402 to 0,7364 | No | ns | >0,9999 |
| 27 | -0,6585 | -2,228 to 0,9108 | No | ns | >0,9999 |
| 28 | 0,008633 | -1,561 to 1,578 | No | ns | >0,9999 |
| 29 | 0,07843 | -1,491 to 1,648 | No | ns | >0,9999 |
| 30 | 0,1215 | -1,448 to 1,691 | No | ns | >0,9999 |
| 31 | 0,1201 | -1,449 to 1,689 | No | ns | >0,9999 |
| 32 | 0,09778 | -1,472 to 1,667 | No | ns | >0,9999 |
| 33 | 0,1022 | -1,467 to 1,671 | No | ns | >0,9999 |
| 34 | 0,09672 | -1,473 to 1,666 | No | ns | >0,9999 |
| 35 | 0,1199 | -1,449 to 1,689 | No | ns | >0,9999 |
| 36 | 0,1115 | -1,458 to 1,681 | No | ns | >0,9999 |
| 37 | 0,1039 | -1,465 to 1,673 | No | ns | >0,9999 |
| 38 | 0,05178 | -1,518 to 1,621 | No | ns | >0,9999 |
| 39 | 0,01918 | -1,550 to 1,588 | No | ns | >0,9999 |
| 40 | 0,01925 | -1,550 to 1,589 | No | ns | >0,9999 |
| 41 | 0,01545 | -1,554 to 1,585 | No | ns | >0,9999 |
| 42 | -0,02265 | -1,592 to 1,547 | No | ns | >0,9999 |
| 43 | 0,001033 | -1,568 to 1,570 | No | ns | >0,9999 |
| 44 | 0,02818 | -1,541 to 1,597 | No | ns | >0,9999 |
| 45 | 0,07722 | -1,492 to 1,647 | No | ns | >0,9999 |
| 46 | 0,1215 | -1,448 to 1,691 | No | ns | >0,9999 |
| 47 | 0,1766 | -1,393 to 1,746 | No | ns | >0,9999 |
| 48 | 0,1580 | -1,411 to 1,727 | No | ns | >0,9999 |
| 49 | 0,1641 | -1,405 to 1,733 | No | ns | >0,9999 |
| 50 | 0,1397 | -1,430 to 1,709 | No | ns | >0,9999 |
| 51 | 0,1227 | -1,447 to 1,692 | No | ns | >0,9999 |
| 52 | 0,1026 | -1,467 to 1,672 | No | ns | >0,9999 |
| 53 | 0,08602 | -1,483 to 1,655 | No | ns | >0,9999 |
| 54 | 0,04920 | -1,520 to 1,619 | No | ns | >0,9999 |
| 55 | 0,05165 | -1,518 to 1,621 | No | ns | >0,9999 |
| 56 | 0,04797 | -1,521 to 1,617 | No | ns | >0,9999 |
| 57 | 0,06490 | -1,504 to 1,634 | No | ns | >0,9999 |
| 58 | 0,06837 | -1,501 to 1,638 | No | ns | >0,9999 |
| 59 | 0,05853 | -1,511 to 1,628 | No | ns | >0,9999 |
| 60 | 0,03623 | -1,533 to 1,606 | No | ns | >0,9999 |
| 61 | 0,05272 | -1,517 to 1,622 | No | ns | >0,9999 |
| 62 | 0,05685 | -1,512 to 1,626 | No | ns | >0,9999 |
| 63 | 0,1154 | -1,454 to 1,685 | No | ns | >0,9999 |
| 64 | 0,1579 | -1,411 to 1,727 | No | ns | >0,9999 |
| 65 | 0,1976 | -1,372 to 1,767 | No | ns | >0,9999 |
| 66 | 0,2034 | -1,366 to 1,773 | No | ns | >0,9999 |
| 67 | 0,3064 | -1,263 to 1,876 | No | ns | >0,9999 |
| 68 | 0,5059 | -1,063 to 2,075 | No | ns | >0,9999 |
| 71 | 1,499 | -0,07048 to 3,068 | No | ns | 0,0935 |
| 72 | 1,663 | 0,09340 to 3,232 | No | ns | 0,207 |
| 73 | 1,384 | -0,1854 to 2,953 | No | ns | 0,2346 |
| 74 | 1,084 | -0,4849 to 2,654 | No | ns | 0,9295 |
| 75 | 1,143 | -0,4268 to 2,712 | No | ns | 0,8280 |
| 76 | 1,106 | -0,4634 to 2,675 | No | ns | 0,8981 |
| 77 | 1,355 | -0,2140 to 2,925 | No | ns | 0,2878 |
| 78 | 1,853 | 0,2833 to 3,422 | Yes | ** | 0,0029 |
| 79 | 1,558 | -0,01083 to 3,128 | No | ns | 0,0552 |
| 80 | 1,443 | -0,1268 to 3,012 | No | ns | 0,1493 |
| 81 | 1,040 | -0,5289 to 2,610 | No | ns | 0,9721 |
| 82 | 0,9768 | -0,5925 to 2,546 | No | ns | 0,9956 |
| 83 | 0,5730 | -0,9963 to 2,142 | No | ns | >0,9999 |
| 84 | 0,2178 | -1,352 to 1,787 | No | ns | >0,9999 |
| 85 | 0,09842 | -1,471 to 1,668 | No | ns | >0,9999 |
| 86 | 0,07445 | -1,495 to 1,644 | No | ns | >0,9999 |
| 87 | 0,07285 | -1,496 to 1,642 | No | ns | >0,9999 |
| 88 | 0,04062 | -1,529 to 1,610 | No | ns | >0,9999 |
| 89 | 0,04402 | -1,525 to 1,613 | No | ns | >0,9999 |
| 90 | 0,06403 | -1,505 to 1,633 | No | ns | >0,9999 |
| 91 | 0,1190 | -1,450 to 1,688 | No | ns | >0,9999 |
| 92 | 0,1667 | -1,403 to 1,736 | No | ns | >0,9999 |
| 93 | 0,2036 | -1,366 to 1,773 | No | ns | >0,9999 |
| 94 | 0,2328 | -1,337 to 1,802 | No | ns | >0,9999 |
| 95 | 0,2515 | -1,318 to 1,821 | No | ns | >0,9999 |
| 96 | 0,2559 | -1,313 to 1,825 | No | ns | >0,9999 |
| 97 | 0,09010 | -1,479 to 1,659 | No | ns | >0,9999 |
| 98 | -0,04817 | -1,617 to 1,521 | No | ns | >0,9999 |
| 99 | -0,04390 | -1,613 to 1,525 | No | ns | >0,9999 |
| 100 | 0,2482 | -1,321 to 1,818 | No | ns | >0,9999 |
| 101 | 0,2960 | -1,273 to 1,865 | No | ns | >0,9999 |
| 102 | 0,2943 | -1,275 to 1,864 | No | ns | >0,9999 |
| 103 | 0,2991 | -1,270 to 1,868 | No | ns | >0,9999 |
| 104 | 0,2596 | -1,310 to 1,829 | No | ns | >0,9999 |
| 105 | 0,2179 | -1,351 to 1,787 | No | ns | >0,9999 |
| 106 | 0,1703 | -1,399 to 1,740 | No | ns | >0,9999 |
| 107 | 0,1356 | -1,434 to 1,705 | No | ns | >0,9999 |
| 108 | 0,1081 | -1,461 to 1,677 | No | ns | >0,9999 |
| 109 | 0,09800 | -1,471 to 1,667 | No | ns | >0,9999 |
| 110 | 0,08167 | -1,488 to 1,651 | No | ns | >0,9999 |
| 111 | 0,1226 | -1,447 to 1,692 | No | ns | >0,9999 |
| 112 | 0,1433 | -1,426 to 1,713 | No | ns | >0,9999 |
| 113 | 0,1239 | -1,445 to 1,693 | No | ns | >0,9999 |
| 114 | 0,1017 | -1,468 to 1,671 | No | ns | >0,9999 |
| 115 | 0,09857 | -1,471 to 1,668 | No | ns | >0,9999 |
| 116 | 0,1169 | -1,452 to 1,686 | No | ns | >0,9999 |
| 117 | 0,1326 | -1,437 to 1,702 | No | ns | >0,9999 |
| 118 | 0,1560 | -1,413 to 1,725 | No | ns | >0,9999 |
| 119 | 0,1954 | -1,374 to 1,765 | No | ns | >0,9999 |
| 120 | 0,2262 | -1,343 to 1,795 | No | ns | >0,9999 |
| 121 | 0,2807 | -1,289 to 1,850 | No | ns | >0,9999 |
| 122 | 0,2631 | -1,306 to 1,832 | No | ns | >0,9999 |
| 123 | 0,2266 | -1,343 to 1,796 | No | ns | >0,9999 |
| 124 | 0,2003 | -1,369 to 1,770 | No | ns | >0,9999 |
| 125 | 0,3811 | -1,188 to 1,950 | No | ns | >0,9999 |
| 126 | 0,3791 | -1,190 to 1,948 | No | ns | >0,9999 |
| 127 | 0,3244 | -1,245 to 1,894 | No | ns | >0,9999 |
| 128 | 0,2923 | -1,277 to 1,862 | No | ns | >0,9999 |
| 129 | 0,2453 | -1,324 to 1,815 | No | ns | >0,9999 |
| 130 | 0,2129 | -1,356 to 1,782 | No | ns | >0,9999 |
| 131 | 0,1849 | -1,384 to 1,754 | No | ns | >0,9999 |
| 132 | 0,1594 | -1,410 to 1,729 | No | ns | >0,9999 |
| 133 | 0,1386 | -1,431 to 1,708 | No | ns | >0,9999 |
| 134 | 0,1484 | -1,421 to 1,718 | No | ns | >0,9999 |
| 135 | 0,1306 | -1,439 to 1,700 | No | ns | >0,9999 |
| 136 | 0,03127 | -1,538 to 1,601 | No | ns | >0,9999 |
| 137 | 0,1496 | -1,420 to 1,719 | No | ns | >0,9999 |
| 138 | 0,2176 | -1,352 to 1,787 | No | ns | >0,9999 |
| 139 | 0,2701 | -1,299 to 1,839 | No | ns | >0,9999 |
| 140 | 0,1339 | -1,435 to 1,703 | No | ns | >0,9999 |
| 141 | 0,09455 | -1,475 to 1,664 | No | ns | >0,9999 |
| 142 | -0,02252 | -1,592 to 1,547 | No | ns | >0,9999 |
| 143 | -0,1161 | -1,685 to 1,453 | No | ns | >0,9999 |
| 145 | 0,5314 | -1,038 to 2,101 | No | ns | >0,9999 |
| 146 | 0,8515 | -0,7178 to 2,421 | No | ns | >0,9999 |
| 147 | 1,087 | -0,4820 to 2,657 | No | ns | 0,9257 |
| 148 | 1,281 | -0,2884 to 2,850 | No | ns | 0,4617 |
| 149 | 0,4073 | -1,162 to 1,977 | No | ns | >0,9999 |
| 150 | 0,2902 | -1,279 to 1,860 | No | ns | >0,9999 |
| 151 | 0,2379 | -1,331 to 1,807 | No | ns | >0,9999 |
| 152 | 0,1059 | -1,463 to 1,675 | No | ns | >0,9999 |
| 153 | 0,04577 | -1,524 to 1,615 | No | ns | >0,9999 |
| 154 | -0,1927 | -1,762 to 1,377 | No | ns | >0,9999 |
| 155 | -0,1416 | -1,711 to 1,428 | No | ns | >0,9999 |
| 156 | -0,1053 | -1,675 to 1,464 | No | ns | >0,9999 |
| 157 | -0,1428 | -1,712 to 1,427 | No | ns | >0,9999 |
| 158 | 0,2225 | -1,347 to 1,792 | No | ns | >0,9999 |
| 159 | 0,4210 | -1,148 to 1,990 | No | ns | >0,9999 |
| 160 | 0,2989 | -1,270 to 1,868 | No | ns | >0,9999 |
| 161 | 0,3435 | -1,226 to 1,913 | No | ns | >0,9999 |
| 162 | 0,4896 | -1,080 to 2,059 | No | ns | >0,9999 |
| 163 | 0,4112 | -1,158 to 1,980 | No | ns | >0,9999 |
| 164 | 0,1765 | -1,393 to 1,746 | No | ns | >0,9999 |
| 165 | 0,02417 | -1,545 to 1,593 | No | ns | >0,9999 |
| 166 | 0,01922 | -1,550 to 1,589 | No | ns | >0,9999 |
| 167 | 0,1414 | -1,428 to 1,711 | No | ns | >0,9999 |
| 168 | 0,1849 | -1,384 to 1,754 | No | ns | >0,9999 |
| 169 | 0,1906 | -1,379 to 1,760 | No | ns | >0,9999 |
| 170 | 0,2266 | -1,343 to 1,796 | No | ns | >0,9999 |
| 171 | 0,2555 | -1,314 to 1,825 | No | ns | >0,9999 |
| 172 | 0,2866 | -1,283 to 1,856 | No | ns | >0,9999 |
| 173 | 0,2782 | -1,291 to 1,848 | No | ns | >0,9999 |
| 174 | 0,2838 | -1,286 to 1,853 | No | ns | >0,9999 |
| 175 | 0,3901 | -1,179 to 1,959 | No | ns | >0,9999 |
| 176 | 0,5244 | -1,045 to 2,094 | No | ns | >0,9999 |
| 177 | 0,5021 | -1,067 to 2,071 | No | ns | >0,9999 |
| 178 | 0,2040 | -1,365 to 1,773 | No | ns | >0,9999 |
| 179 | 0,1525 | -1,417 to 1,722 | No | ns | >0,9999 |
| 180 | 0,07663 | -1,493 to 1,646 | No | ns | >0,9999 |
| 181 | 0,1202 | -1,449 to 1,689 | No | ns | >0,9999 |
| 182 | -0,06235 | -1,632 to 1,507 | No | ns | >0,9999 |
| 183 | -0,1550 | -1,724 to 1,414 | No | ns | >0,9999 |
| 184 | 0,1200 | -1,449 to 1,689 | No | ns | >0,9999 |
| 185 | 0,3601 | -1,209 to 1,929 | No | ns | >0,9999 |
| 186 | 0,2735 | -1,296 to 1,843 | No | ns | >0,9999 |
| 187 | 0,06557 | -1,504 to 1,635 | No | ns | >0,9999 |
| 188 | -0,006367 | -1,576 to 1,563 | No | ns | >0,9999 |
| 189 | 0,1482 | -1,421 to 1,717 | No | ns | >0,9999 |
| 190 | 0,1683 | -1,401 to 1,738 | No | ns | >0,9999 |
| 191 | 0,2178 | -1,351 to 1,787 | No | ns | >0,9999 |
| 192 | 0,2063 | -1,363 to 1,776 | No | ns | >0,9999 |
| 193 | 0,1819 | -1,387 to 1,751 | No | ns | >0,9999 |
| 194 | 0,1470 | -1,422 to 1,716 | No | ns | >0,9999 |
| 195 | 0,1175 | -1,452 to 1,687 | No | ns | >0,9999 |
| 196 | 0,08248 | -1,487 to 1,652 | No | ns | >0,9999 |
| 197 | 0,05312 | -1,516 to 1,622 | No | ns | >0,9999 |
| 198 | -0,02153 | -1,591 to 1,548 | No | ns | >0,9999 |
| 199 | -0,02647 | -1,596 to 1,543 | No | ns | >0,9999 |
| 200 | -0,05638 | -1,626 to 1,513 | No | ns | >0,9999 |
| 201 | 0,01135 | -1,558 to 1,581 | No | ns | >0,9999 |
| 202 | 0,02795 | -1,541 to 1,597 | No | ns | >0,9999 |
| 203 | 0,06480 | -1,505 to 1,634 | No | ns | >0,9999 |
| 204 | 0,08737 | -1,482 to 1,657 | No | ns | >0,9999 |
| 205 | 0,1054 | -1,464 to 1,675 | No | ns | >0,9999 |
| 206 | 0,1315 | -1,438 to 1,701 | No | ns | >0,9999 |
| 207 | 0,1754 | -1,394 to 1,745 | No | ns | >0,9999 |
| 208 | 0,2154 | -1,354 to 1,785 | No | ns | >0,9999 |
| 209 | 0,2319 | -1,337 to 1,801 | No | ns | >0,9999 |
| 210 | 0,2664 | -1,303 to 1,836 | No | ns | >0,9999 |
| 211 | 0,1726 | -1,397 to 1,742 | No | ns | >0,9999 |
| 212 | 0,2060 | -1,363 to 1,775 | No | ns | >0,9999 |
| 213 | -0,009050 | -1,578 to 1,560 | No | ns | >0,9999 |
| 214 | 0,1831 | -1,386 to 1,752 | No | ns | >0,9999 |
| 215 | 0,1034 | -1,466 to 1,673 | No | ns | >0,9999 |
| 216 | 0,09808 | -1,471 to 1,667 | No | ns | >0,9999 |
| 217 | 0,1122 | -1,457 to 1,681 | No | ns | >0,9999 |
| 218 | 0,07728 | -1,492 to 1,647 | No | ns | >0,9999 |
| 219 | 0,04660 | -1,523 to 1,616 | No | ns | >0,9999 |
| 220 | 0,04433 | -1,525 to 1,614 | No | ns | >0,9999 |
| 221 | 0,06770 | -1,502 to 1,637 | No | ns | >0,9999 |
| 222 | 0,1077 | -1,462 to 1,677 | No | ns | >0,9999 |
| 223 | 0,1143 | -1,455 to 1,684 | No | ns | >0,9999 |
| 224 | 0,1089 | -1,460 to 1,678 | No | ns | >0,9999 |
| 225 | 0,07860 | -1,491 to 1,648 | No | ns | >0,9999 |
| 226 | 0,04960 | -1,520 to 1,619 | No | ns | >0,9999 |
| 227 | 0,03652 | -1,533 to 1,606 | No | ns | >0,9999 |
| 228 | 0,04048 | -1,529 to 1,610 | No | ns | >0,9999 |
| 229 | 0,04085 | -1,528 to 1,610 | No | ns | >0,9999 |
| 230 | 0,04428 | -1,525 to 1,614 | No | ns | >0,9999 |
| 231 | 0,04500 | -1,524 to 1,614 | No | ns | >0,9999 |
| 232 | 0,01142 | -1,558 to 1,581 | No | ns | >0,9999 |
| 233 | -0,04212 | -1,611 to 1,527 | No | ns | >0,9999 |
| 234 | -0,06228 | -1,632 to 1,507 | No | ns | >0,9999 |
| 235 | -0,06130 | -1,631 to 1,508 | No | ns | >0,9999 |
| 236 | 0,03678 | -1,533 to 1,606 | No | ns | >0,9999 |
| 237 | 0,06193 | -1,507 to 1,631 | No | ns | >0,9999 |
| 238 | 0,08870 | -1,481 to 1,658 | No | ns | >0,9999 |
| 239 | 0,1230 | -1,446 to 1,692 | No | ns | >0,9999 |
| 240 | 0,1599 | -1,409 to 1,729 | No | ns | >0,9999 |
| 241 | 0,1866 | -1,383 to 1,756 | No | ns | >0,9999 |
| 242 | 0,2026 | -1,367 to 1,772 | No | ns | >0,9999 |
| 243 | 0,1846 | -1,385 to 1,754 | No | ns | >0,9999 |
| 244 | 0,1943 | -1,375 to 1,764 | No | ns | >0,9999 |
| 245 | 0,2541 | -1,315 to 1,823 | No | ns | >0,9999 |
| 246 | 0,3629 | -1,206 to 1,932 | No | ns | >0,9999 |
| 247 | 0,2525 | -1,317 to 1,822 | No | ns | >0,9999 |
| 248 | 0,8441 | -0,7252 to 2,413 | No | ns | >0,9999 |
| 249 | 1,800 | 0,2307 to 3,369 | Yes | ** | 0,0051 |
| 250 | 1,981 | 0,4121 to 3,551 | Yes | *** | 0,0007 |
| 251 | 1,504 | -0,06485 to 3,074 | No | ns | 0,0890 |
| 252 | 1,468 | -0,1018 to 3,037 | No | ns | 0,1217 |
| 253 | 1,063 | -0,5060 to 2,633 | No | ns | 0,9534 |
| 254 | -0,1437 | -1,713 to 1,426 | No | ns | >0,9999 |
| 255 | -0,2500 | -1,819 to 1,319 | No | ns | >0,9999 |
| 256 | 0,03272 | -1,537 to 1,602 | No | ns | >0,9999 |
| 257 | -0,3874 | -1,957 to 1,182 | No | ns | >0,9999 |
| 258 | -0,4291 | -1,998 to 1,140 | No | ns | >0,9999 |
| 259 | -0,04410 | -1,613 to 1,525 | No | ns | >0,9999 |
| 260 | 0,5205 | -1,049 to 2,090 | No | ns | >0,9999 |
| 261 | 0,4462 | -1,123 to 2,016 | No | ns | >0,9999 |
| 262 | 0,2288 | -1,340 to 1,798 | No | ns | >0,9999 |
| 263 | 0,2676 | -1,302 to 1,837 | No | ns | >0,9999 |
| 264 | 0,1012 | -1,468 to 1,670 | No | ns | >0,9999 |
| 265 | 0,1285 | -1,441 to 1,698 | No | ns | >0,9999 |
| 266 | 0,1797 | -1,390 to 1,749 | No | ns | >0,9999 |
| 267 | 0,2085 | -1,361 to 1,778 | No | ns | >0,9999 |
| 268 | 0,1588 | -1,411 to 1,728 | No | ns | >0,9999 |
| 269 | 0,1300 | -1,439 to 1,699 | No | ns | >0,9999 |
| 270 | 0,09563 | -1,474 to 1,665 | No | ns | >0,9999 |
| 271 | 0,08532 | -1,484 to 1,655 | No | ns | >0,9999 |
| 272 | 0,05353 | -1,516 to 1,623 | No | ns | >0,9999 |
| 273 | 0,05000 | -1,519 to 1,619 | No | ns | >0,9999 |
| 274 | 0,06108 | -1,508 to 1,630 | No | ns | >0,9999 |
| 275 | 0,06670 | -1,503 to 1,636 | No | ns | >0,9999 |
| 276 | 0,07000 | -1,499 to 1,639 | No | ns | >0,9999 |
| 277 | 0,08763 | -1,482 to 1,657 | No | ns | >0,9999 |
| 278 | 0,09057 | -1,479 to 1,660 | No | ns | >0,9999 |
| 279 | 0,1078 | -1,461 to 1,677 | No | ns | >0,9999 |
| 280 | 0,1234 | -1,446 to 1,693 | No | ns | >0,9999 |
| 281 | 0,1413 | -1,428 to 1,711 | No | ns | >0,9999 |
| 282 | 0,1514 | -1,418 to 1,721 | No | ns | >0,9999 |
| 283 | 0,1191 | -1,450 to 1,688 | No | ns | >0,9999 |
| 284 | 0,07558 | -1,494 to 1,645 | No | ns | >0,9999 |
| 285 | 0,07423 | -1,495 to 1,644 | No | ns | >0,9999 |
| 286 | 0,07518 | -1,494 to 1,644 | No | ns | >0,9999 |
| 287 | 0,1054 | -1,464 to 1,675 | No | ns | >0,9999 |
| 288 | 0,1002 | -1,469 to 1,670 | No | ns | >0,9999 |
| 289 | 0,08975 | -1,480 to 1,659 | No | ns | >0,9999 |
| 290 | 0,07340 | -1,496 to 1,643 | No | ns | >0,9999 |
| 291 | 0,05165 | -1,518 to 1,621 | No | ns | >0,9999 |
| 292 | 0,04182 | -1,527 to 1,611 | No | ns | >0,9999 |
| 293 | 0,01907 | -1,550 to 1,588 | No | ns | >0,9999 |
| 294 | 0,004067 | -1,565 to 1,573 | No | ns | >0,9999 |
| 295 | -0,01487 | -1,584 to 1,554 | No | ns | >0,9999 |
| 296 | -0,02013 | -1,589 to 1,549 | No | ns | >0,9999 |

**Tab.2:** Statistical analysis of rmsf plots from wt and B.1.1.7 spike protein residues 800-900 (n =6; two-way ANOVA; significance assumed for *p<0.05).

| Compare each cell mean with the other cell mean in that row |  |  |  |  |  |
| --- | --- | --- | --- | --- | --- |
| Number of families | 1 |  |  |  |  |
| Number of comparisons per family | 100 |  |  |  |  |
| Alpha | 0,05 |  |  |  |  |
| Sidak's multiple comparisons test | Mean Diff, | 95,00% CI of diff, | Significant? | Summary | Adjusted P Value |
| residue number |  |  |  |  |  |
| 800 | -0,1280 | -1,995 to 1,739 | No | ns | >0,9999 |
| 801 | -0,2166 | -2,084 to 1,650 | No | ns | >0,9999 |
| 802 | -0,2190 | -2,086 to 1,648 | No | ns | >0,9999 |
| 803 | -0,2493 | -2,116 to 1,618 | No | ns | >0,9999 |
| 804 | -0,3069 | -2,174 to 1,560 | No | ns | >0,9999 |
| 805 | -0,2610 | -2,128 to 1,606 | No | ns | >0,9999 |
| 806 | -0,2261 | -2,093 to 1,641 | No | ns | >0,9999 |
| 807 | -0,1018 | -1,969 to 1,765 | No | ns | >0,9999 |
| 808 | 0,06743 | -1,799 to 1,934 | No | ns | >0,9999 |
| 809 | -0,04395 | -1,911 to 1,823 | No | ns | >0,9999 |
| 810 | -0,1864 | -2,053 to 1,681 | No | ns | >0,9999 |
| 811 | -0,2875 | -2,154 to 1,579 | No | ns | >0,9999 |
| 812 | -0,3187 | -2,186 to 1,548 | No | ns | >0,9999 |
| 813 | -0,4475 | -2,314 to 1,419 | No | ns | >0,9999 |
| 814 | -0,3771 | -2,244 to 1,490 | No | ns | >0,9999 |
| 815 | -0,3110 | -2,178 to 1,556 | No | ns | >0,9999 |
| 816 | -0,2772 | -2,144 to 1,590 | No | ns | >0,9999 |
| 817 | -0,2562 | -2,123 to 1,611 | No | ns | >0,9999 |
| 818 | -0,2571 | -2,124 to 1,610 | No | ns | >0,9999 |
| 819 | -0,2522 | -2,119 to 1,615 | No | ns | >0,9999 |
| 820 | -0,2137 | -2,081 to 1,653 | No | ns | >0,9999 |
| 821 | -0,2046 | -2,071 to 1,662 | No | ns | >0,9999 |
| 822 | -0,1648 | -2,032 to 1,702 | No | ns | >0,9999 |
| 823 | -0,2029 | -2,070 to 1,664 | No | ns | >0,9999 |
| 824 | -0,2830 | -2,150 to 1,584 | No | ns | >0,9999 |
| 825 | -0,3213 | -2,188 to 1,546 | No | ns | >0,9999 |
| 826 | -0,1972 | -2,064 to 1,670 | No | ns | >0,9999 |
| 827 | -0,3541 | -2,221 to 1,513 | No | ns | >0,9999 |
| 828 | -0,4093 | -2,276 to 1,458 | No | ns | >0,9999 |
| 829 | -0,9103 | -2,777 to 0,9567 | No | ns | >0,9999 |
| 830 | -1,089 | -2,956 to 0,7779 | No | ns | 0,9862 |
| 831 | -1,212 | -3,079 to 0,6549 | No | ns | 0,9084 |
| 832 | -1,656 | -3,522 to 0,2113 | No | ns | 0,1845 |
| 833 | -1,782 | -3,649 to 0,08476 | No | ns | 0,0867 |
| 834 | -1,768 | -3,635 to<br>0,09891 | No | ns | 0,0947 |
| 835 | -2,051 | -3,918 to -0,1844 | Yes | * | 0,0137 |
| 836 | -2,291 | -4,158 to -0,4240 | Yes | ** | 0,0021 |
| 837 | -2,393 | -4,260 to -0,5262 | Yes | *** | 0,0009 |
| 838 | -2,537 | -4,404 to -0,6704 | Yes | *** | 0,0003 |
| 839 | -1,878 | -3,745 to -<br>0,01122 | Yes | * | 0,0464 |
| 840 | -2,151 | -4,017 to -0,2836 | Yes | ** | 0,0065 |
| 841 | -2,035 | -3,902 to -0,1686 | Yes | * | 0,0154 |
| 842 | -2,393 | -4,260 to -0,5264 | Yes | *** | 0,0009 |
| 843 | -2,179 | -4,046 to -0,3123 | Yes | ** | 0,0052 |
| 844 | -1,754 | -3,620 to 0,1133 | No | ns | 0,1036 |
| 845 | -1,753 | -3,620 to 0,1140 | No | ns | 0,1040 |
| 846 | -1,470 | -3,337 to 0,3967 | No | ns | 0,4579 |
| 847 | -1,228 | -3,095 to 0,6391 | No | ns | 0,8905 |
| 848 | -1,048 | -2,915 to 0,8192 | No | ns | 0,9942 |
| 849 | -0,9947 | -2,862 to 0,8722 | No | ns | 0,9985 |
| 850 | -0,7876 | -2,654 to 1,079 | No | ns | >0,9999 |
| 851 | -1,005 | -2,872 to 0,8620 | No | ns | 0,9980 |
| 852 | -0,8561 | -2,723 to 1,011 | No | ns | >0,9999 |
| 853 | -0,5983 | -2,465 to 1,269 | No | ns | >0,9999 |
| 854 | -0,3093 | -2,176 to 1,558 | No | ns | >0,9999 |
| 855 | -0,04725 | -1,914 to 1,820 | No | ns | >0,9999 |
| 856 | -0,07502 | -1,942 to 1,792 | No | ns | >0,9999 |
| 857 | -0,1309 | -1,998 to 1,736 | No | ns | >0,9999 |
| 858 | -0,1545 | -2,021 to 1,712 | No | ns | >0,9999 |
| 859 | -0,2139 | -2,081 to 1,653 | No | ns | >0,9999 |
| 860 | -0,2246 | -2,091 to 1,642 | No | ns | >0,9999 |
| 861 | -0,2097 | -2,077 to 1,657 | No | ns | >0,9999 |
| 862 | -0,2924 | -2,159 to 1,575 | No | ns | >0,9999 |
| 863 | -0,3368 | -2,204 to 1,530 | No | ns | >0,9999 |
| 864 | -0,3380 | -2,205 to 1,529 | No | ns | >0,9999 |
| 865 | -0,3322 | -2,199 to 1,535 | No | ns | >0,9999 |
| 866 | -0,2914 | -2,158 to 1,576 | No | ns | >0,9999 |
| 867 | -0,3157 | -2,183 to 1,551 | No | ns | >0,9999 |
| 868 | -0,3822 | -2,249 to 1,485 | No | ns | >0,9999 |
| 869 | -0,3861 | -2,253 to 1,481 | No | ns | >0,9999 |
| 870 | -0,3807 | -2,248 to 1,486 | No | ns | >0,9999 |
| 871 | -0,3803 | -2,247 to 1,487 | No | ns | >0,9999 |
| 872 | -0,3900 | -2,257 to 1,477 | No | ns | >0,9999 |
| 873 | -0,4014 | -2,268 to 1,466 | No | ns | >0,9999 |
| 874 | -0,3739 | -2,241 to 1,493 | No | ns | >0,9999 |
| 875 | -0,3471 | -2,214 to 1,520 | No | ns | >0,9999 |
| 876 | -0,3396 | -2,206 to 1,527 | No | ns | >0,9999 |
| 877 | -0,3380 | -2,205 to 1,529 | No | ns | >0,9999 |
| 878 | -0,3264 | -2,193 to 1,540 | No | ns | >0,9999 |
| 879 | -0,2904 | -2,157 to 1,576 | No | ns | >0,9999 |
| 880 | -0,2508 | -2,118 to 1,616 | No | ns | >0,9999 |
| 881 | -0,2283 | -2,095 to 1,639 | No | ns | >0,9999 |
| 882 | -0,1984 | -2,065 to 1,669 | No | ns | >0,9999 |
| 883 | -0,1742 | -2,041 to 1,693 | No | ns | >0,9999 |
| 884 | -0,1965 | -2,063 to 1,670 | No | ns | >0,9999 |
| 885 | -0,08338 | -1,950 to 1,784 | No | ns | >0,9999 |
| 886 | -0,2021 | -2,069 to 1,665 | No | ns | >0,9999 |
| 887 | -0,3500 | -2,217 to 1,517 | No | ns | >0,9999 |
| 888 | -0,4662 | -2,333 to 1,401 | No | ns | >0,9999 |
| 889 | -0,5197 | -2,387 to 1,347 | No | ns | >0,9999 |
| 890 | -0,3981 | -2,265 to 1,469 | No | ns | >0,9999 |
| 891 | -0,3144 | -2,181 to 1,552 | No | ns | >0,9999 |
| 892 | -0,1745 | -2,041 to 1,692 | No | ns | >0,9999 |
| 893 | -0,1717 | -2,039 to 1,695 | No | ns | >0,9999 |
| 894 | -0,1517 | -2,019 to 1,715 | No | ns | >0,9999 |
| 895 | -0,1323 | -1,999 to 1,735 | No | ns | >0,9999 |
| 896 | -0,1209 | -1,988 to 1,746 | No | ns | >0,9999 |
| 897 | -0,1356 | -2,002 to 1,731 | No | ns | >0,9999 |
| 898 | -0,1097 | -1,977 to 1,757 | No | ns | >0,9999 |
| 899 | -0,09742 | -1,964 to 1,769 | No | ns | >0,9999 |

### D614G induces flexibility around the fusion peptide via salt bridge rearrangement

We found lysine 835 and lysine 854 as positively charged amino acid residues in the region of this increased flexibility. Analyzing the wild type structure during MD simulation revealed interchain salt bridge formation between lysine 835 from one chain and aspartate 568 from the neighboring chain and interchain salt bridge formation between lysine 854 and aspartate 614 (Fig2 a). We also identified interchain salt bridges between arginine 646 and glutamate 868 and aspartate 867 (Fig2 a) although to a lesser extent than the other two described charged residue pairs. To analyze the consistency of these identified salt bridges we measured the interatomic distance over all three trimer subunits for both simulation runs. Therefore, we defined distances between the carbon atoms of the carboxyl groups of aspartate or glutamate and the nitrogen of the NH_3_^+^ group in the side chain of lysine or the carbon atom in ζ-position (most distal carbon in side chain) in arginine under 4 Å (preferred distance for salt bridge formation), between 4-5 Å (important for arginine residues due to the usage of the carbon atom in ζ-position) and above 5 Å (for no salt bridge formation). The interchain salt bridge, which can only be present in wild type, formed by aspartate 614 and lysine 854, is in the preferred distance of under 4 Å in about 60% of the simulation time. This indicates a certain importance for this salt bridge for local stability. For the salt bridge formed by lysine 835 from one subunit and aspartate 568 from an adjacent subunit we found distances smaller 4 Å in over 40% of the time for the wt S protein (Fig2 a). Additionally, there was no interaction found between lysine 854 and aspartate 586, arguing for a stable conformation of this fusion peptide adjacent loop region in wild type as it is supported by two salt bridges. Analyzing the same salt bridges in the B.1.1.7 variant reveals a different salt bridge pattern. As aspartate 614 is missing (D614G), lysine 854 is unmatched in this variant. We found that both lysine residues in this region, lysine 854 and lysine 835 form partial salt bridges with aspartate 568. However, even combined, both residues are only in the preferred distance of under 4 Å in about 20% of the time. To analyze the direct effect of the D614G mutation in this matter, we also calculated MD simulations of a wt variant with an inserted glycine for aspartate at position 614 (wt+D614G). Calculating the RMSF values also revealed an increase in flexibility in the region of lysine 854 (sFig3 a). Analyzing the distances between lysine residues 854 and 835 with aspartate 568 also revealed a partial interaction between lysine 854 and aspartate 568 as found in the B.1.1.7 variant (sFig3 b). However, the interaction between lysine 835 and aspartate 568 is as strong as in wt, indicating an additional destabilizing mechanism in the B.1.1.7 variant. The additional destabilization in B.1.1.7 could be explained by the A570D mutation (Fig2 a, orange box). The newly inserted aspartate engages with lysine 964 in about 20% of the time in a preferred salt bridge distance of under 4 Å and could thereby destabilize interactions of aspartate 568 (Fig2 a). For all three variants, wt, wt+D614G and B.1.1.7 partial interaction between arginine 646 with aspartate 867 or glutamate 868 was found (Fig2 a, sFig3 b). As these residues are in close proximity of arginine 815, that is part of the S2’ cleavage site, we also analyzed interactions of this residue with surrounding negatively charged residues (Fig2 c) and found interactions with aspartate 820 and to a minor extend with aspartate 867 in this pre-cleavage state (Fig2 d). To analyze the fate of arginine 815 upon cleavage by e.g. TMPRSS2, we performed *in silico* cleavage between arginine 815 and serine 816 and simulated wt and B.1.1.7. The RMSF values showed the known pattern of increased flexibility for B.1.1.7 (Fig2 d) and we also analyzed salt bridges (Fig2 e). In contrast to the pre-cleavage state, where arginine 815 is in contact with aspartate 820 this interaction is lost in the post-cleavage state and arginine 815 engages with glutamate 819 in wt and B.1.1.7. In wt an additional salt bridge forms in the post-cleavage state between arginine 815 and glutamate 868 the interaction with aspartate 867 remains similar (Fig2 f). In contrast, arginine 815 from B.1.1.7 engages mainly with aspartate 867 in the post-cleavage state and there is only a small interaction with glutamate 868 (Fig2 f). These salt bridge interactions indicating a non-covalent stabilization after proteolytic priming at the S2’ position that requires a certain conformational rearrangement in this area. We also re-analyzed the stabilizing salt bridges between aspartate 614 and lysine 854 and aspartate 568 and lysine 835 in wt (Fig2 g) and found almost no change as compared to pre-cleavage conditions (Fig2 a). For B.1.1.7 we found, that the small interaction between aspartate 568 and lysine 835 is completely lost in the post-cleavage state and only the small interaction between aspartate 568 and lysine 854 remains in B.1.1.7 (Fig2 g). Of note, we also identified that interactions between arginine 646 and aspartate 867 and glutamate 868 are different in wt and B.1.1.7. While there is a more even distribution of contact formation between arginine 646 and the two negatively charged residues in wt, there is a clear preference towards glutamate 868 in B.1.1.7 (Fig2 g). Taken together, we identified how the D614 mutation (in combination with the A570D mutation) induces a loss of conformational stability that increases flexibility in a pre- and post-cleavage state. We also identified changes in salt bridge engagement of arginine 815 between a pre-cleavage (continuous protein chain) and post-cleavage state (discontinuous protein chain), that then stabilizes the new C-terminus via salt bridge rearrangement. To analyze whether this increase in flexibility in B.1.1.7 is also propagated towards the RBD over the entire S protein or is a local phenomenon, we analyzed the RMSD values over the simulation time for the RBDs (sFig3 c-e). The RMSD values were low and very similar for the variants analyzed (wt, wt+D614G and B.1.1.7). Thus we hypothesized that changes in RBD behavior are a direct effect of the N501Y mutation and further analyzed it.

**Figure 2:**
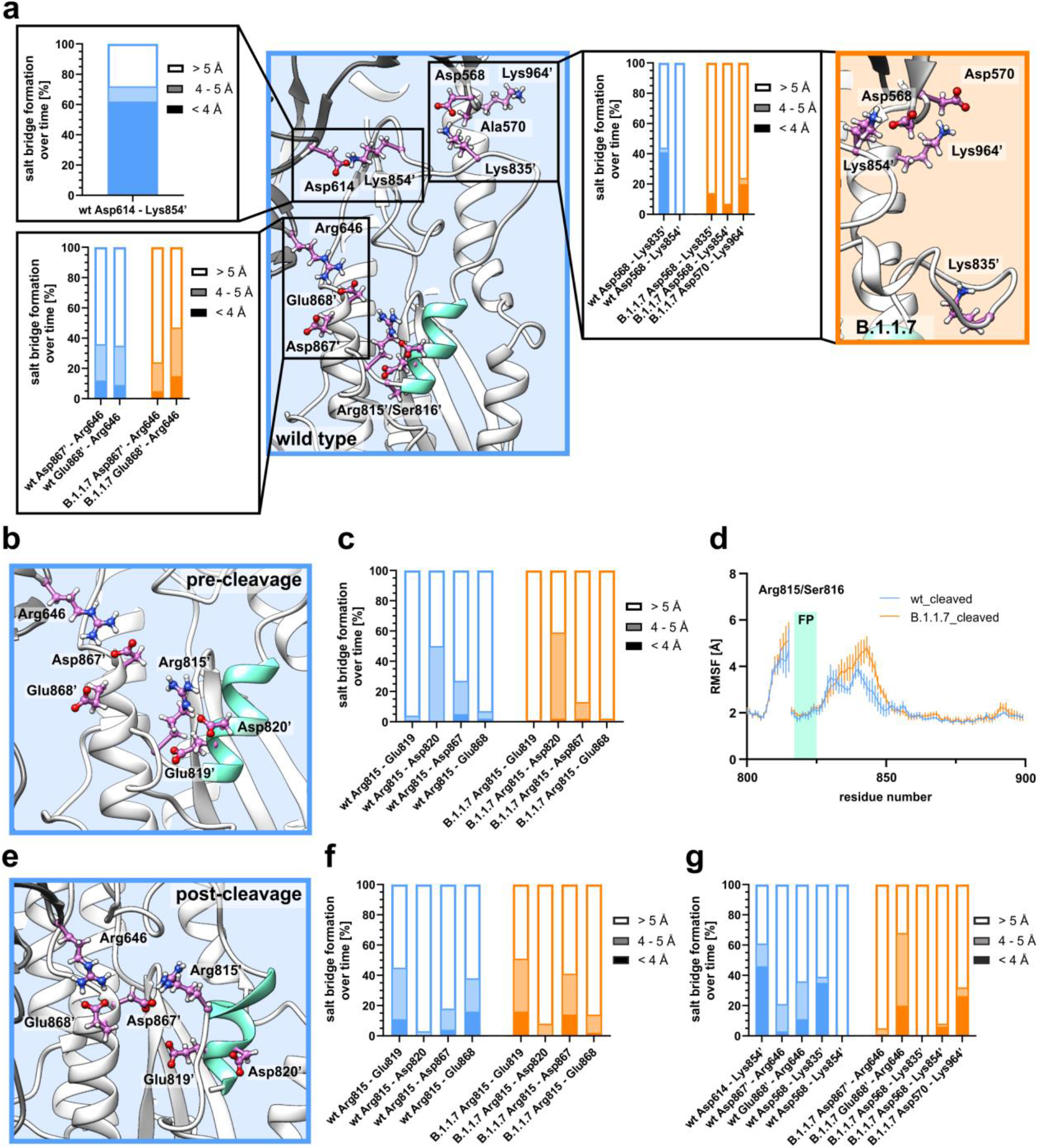
Salt bridge stability is altered upon D614 mutation in pre- and post-cleavage state. a) Stability of a fusion peptide (aquamarine) adjacent loop region is mediated by salt bridge formation between aspartate 614 (Asp614) from one chain and lysine 854 (Lys854’; ‘ denotes residues from a neighboring chain) from the neighboring chain in wild type (wt; blue background). An additional salt bridge is formed by aspartate 568 (Asp568) and lysine 835 (Lys835’) from the neighboring chain in wt. In B.1.1.7 (orange background) this interaction is weaker and the newly inserted aspartate at position 570 (A570D) forms an additional salt bridge with lysine 964 (Lys964’). In close proximity of arginine 815 (part of the S2’ cleavage site) ionic interaction was measured between arginine 646 (Arg646) and aspartate 867 (Asp867’) and glutamate 868 (Glu868’). b) Structural representation of the wt pre-cleavage state (continuous polypeptide chain; similar in B.1.1.7) with negatively charged residues glutamate 819 (Glu819), aspartate 820 (Asp820), aspartate 867 (Asp867) and glutamate 868 (Glu868) shown around arginine 815 (Arg815). c) Percentage of salt bridge formation over time for four negatively charged residues with arginine 815 in the pre-cleavage state. d) Average RMSF values plotted against the residue numbers for wt and B.1.1.7 after *in silico* proteolytic cleavage at the S2’ site (aquamarine bar; FP = fusion peptide). e) Structural representation of the wt post-cleavage state (discontinuous polypeptide chain with a break between arginine 815 and serine 816; similar in B.1.1.7) with negatively charged residues glutamate 819 (Glu819), aspartate 820 (Asp820), aspartate 867 (Asp867) and glutamate 868 (Glu868) shown around arginine 815 (Arg815). f) Percentage of salt bridge formation over time for four different residue pairs in the post-cleavage state. g) Percentage of salt bridge formation over time for the stabilizing salt bridge pairs as analyzed in a).

### N501Y replacement displaces glutamine 498 from the SARS-CoV-2 ACE2 interface

In order to analyze binding of the S protein to the ACE2 receptor we used the isolated receptor-binding domain (RBD) in complex with ACE2 (PDB ID code: 7KMB^16^) as starting structure for molecular dynamics simulations. The only amino acid exchange localized into the RBD is the N501Y exchange that resides at the RBD-ACE2 interface (sFig1 a). Simulation for 500 ns revealed no marked differences in structural flexibility between wt and B.1.1.7, as it can be deduced from the RMSF values (sFig4 a,b). Compared to the MD simulation of the trimeric S protein without ACE2 (unbound state), the overall flexibility was very similar, but the bound RBD (RBD-ACE2 complex) showed markedly lower RMSF values in the region between RBD residue 475 and residue 520, where most of the RBD residues are located which closely interact with ACE2 (sFig4 c). To identify changes between wt and B.1.1.7 we analyzed contacts below 5 Å over time. For the B.1.1.7 variant we identified that glutamine 498 loses almost all contacts to ACE2 residues aspartate 38 and lysine 353, while the number of contacts is strongly reduced for tyrosine 41 (sFig5 a, Tab. 3). We also identified a markedly increased number of contacts for the newly inserted tyrosine at position 501 to lysine 353 in B.1.1.7 when compared to the wt asparagine residue (sFig5 a, individual runs shown in sFig6 and sFig7). Structural analysis revealed the expulsion of glutamine 498 from the RBD-ACE2 interface due to the bulkier tyrosine side chain in the B.1.1.7 variant (sFig5 b,c). We also noticed a change in side chain conformation for lysine 353 explaining the altered contact numbers (sFig5 b,c). We were now interested in individual binding energies and decomposed the RBD-ACE2 interface.

**Tab.3:**
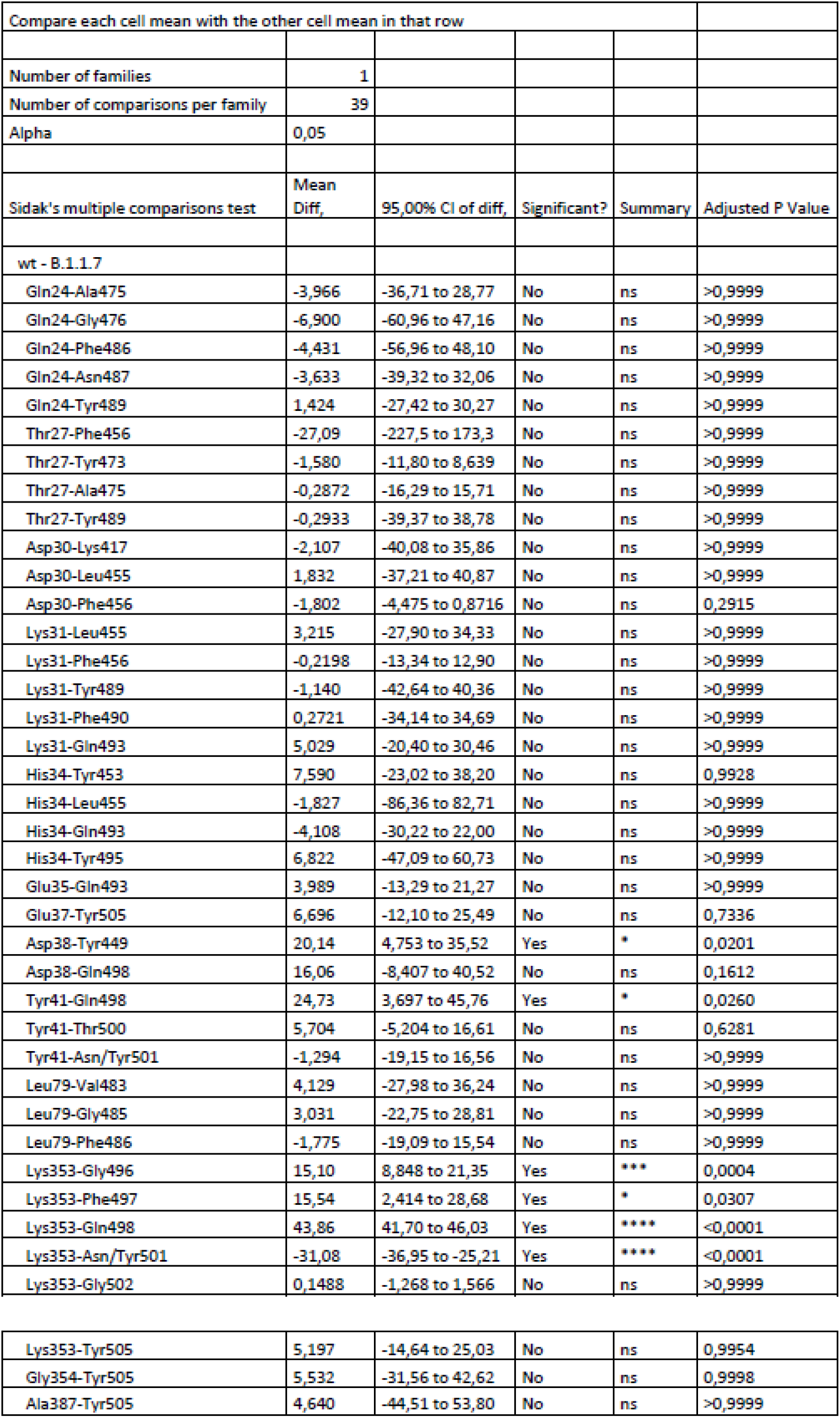
Statistical analysis of contacts between residues from the receptor binding domain and ACE2 from wt and B.1.1.7 (n = 4; two-way ANOVA; significance assumed for *p<0.05).

| Compare each cell mean with the other cell mean in that row |  |  |  |  |  |
| --- | --- | --- | --- | --- | --- |
| Number of families | 1 |  |  |  |  |
| Number of comparisons per family | 39 |  |  |  |  |
| Alpha | 0,05 |  |  |  |  |
|  | Mean |  |  |  |  |
| Sidak's multiple comparisons test | Diff, | 95,00% CI of diff, | Significant? | Summary | Adjusted P Value |
| wt - B.1.1.7 |  |  |  |  |  |
| Gln24-Ala475 | -3,966 | -36,71 to 28,77 | No | ns | >0,9999 |
| Gln24-Gly476 | -6,900 | -60,96 to 47,16 | No | ns | >0,9999 |
| Gln24-Phe486 | -4,431 | -56,96 to 48,10 | No | ns | >0,9999 |
| Gln24-Asn487 | -3,633 | -39,32 to 32,06 | No | ns | >0,9999 |
| Gln24-Tyr489 | 1,424 | -27,42 to 30,27 | No | ns | >0,9999 |
| Thr27-Phe456 | -27,09 | -227,5 to 173,3 | No | ns | >0,9999 |
| Thr27-Tyr473 | -1,580 | -11,80 to 8,639 | No | ns | >0,9999 |
| Thr27-Ala475 | -0,2872 | -16,29 to 15,71 | No | ns | >0,9999 |
| Thr27-Tyr489 | -0,2933 | -39,37 to 38,78 | No | ns | >0,9999 |
| Asp30-Lys417 | -2,107 | -40,08 to 35,86 | No | ns | >0,9999 |
| Asp30-Leu455 | 1,832 | -37,21 to 40,87 | No | ns | >0,9999 |
| Asp30-Phe456 | -1,802 | -4,475 to 0,8716 | No | ns | 0,2915 |
| Lys31-Leu455 | 3,215 | -27,90 to 34,33 | No | ns | >0,9999 |
| Lys31-Phe456 | -0,2198 | -13,34 to 12,90 | No | ns | >0,9999 |
| Lys31-Tyr489 | -1,140 | -42,64 to 40,36 | No | ns | >0,9999 |
| Lys31-Phe490 | 0,2721 | -34,14 to 34,69 | No | ns | >0,9999 |
| Lys31-Gln493 | 5,029 | -20,40 to 30,46 | No | ns | >0,9999 |
| His34-Tyr453 | 7,590 | -23,02 to 38,20 | No | ns | 0,9928 |
| His34-Leu455 | -1,827 | -86,36 to 82,71 | No | ns | >0,9999 |
| His34-Gln493 | -4,108 | -30,22 to 22,00 | No | ns | >0,9999 |
| His34-Tyr495 | 6,822 | -47,09 to 60,73 | No | ns | >0,9999 |
| Glu35-Gln493 | 3,989 | -13,29 to 21,27 | No | ns | >0,9999 |
| Glu37-Tyr505 | 6,696 | -12,10 to 25,49 | No | ns | 0,7336 |
| Asp38-Tyr449 | 20,14 | 4,753 to 35,52 | Yes | * | 0,0201 |
| Asp38-Gln498 | 16,06 | -8,407 to 40,52 | No | ns | 0,1612 |
| Tyr41-Gln498 | 24,73 | 3,697 to 45,76 | Yes | * | 0,0260 |
| Tyr41-Thr500 | 5,704 | -5,204 to 16,61 | No | ns | 0,6281 |
| Tyr41-Asn/Tyr501 | -1,294 | -19,15 to 16,56 | No | ns | >0,9999 |
| Leu79-Val483 | 4,129 | -27,98 to 36,24 | No | ns | >0,9999 |
| Leu79-Gly485 | 3,031 | -22,75 to 28,81 | No | ns | >0,9999 |
| Leu79-Phe486 | -1,775 | -19,09 to 15,54 | No | ns | >0,9999 |
| Lys353-Gly496 | 15,10 | 8,848 to 21,35 | Yes | *** | 0,0004 |
| Lys353-Phe497 | 15,54 | 2,414 to 28,68 | Yes | * | 0,0307 |
| Lys353-Gln498 | 43,86 | 41,70 to 46,03 | Yes | **** | <0,0001 |
| Lys353-Asn/Tyr501 | -31,08 | -36,95 to -25,21 | Yes | **** | <0,0001 |
| Lys353-Gly502 | 0,1488 | -1,268 to 1,566 | No | ns | >0,9999 |
| Lys353-Tyr505 | 5,197 | -14,64 to 25,03 | No | ns | 0,9954 |
| Gly354-Tyr505 | 5,532 | -31,56 to 42,62 | No | ns | 0,9998 |
| Ala387-Tyr505 | 4,640 | -44,51 to 53,80 | No | ns | >0,9999 |

### The N501Y mutation lowers electrostatic binding in B.1.1.7

To decompose the electrostatic linear interaction energy all residues within 4 Å of ACE2 were analyzed. We identified the main electrostatic interaction in the central part of the RBD-ACE2 interface for wt and B.1.1.7. The electrostatic interaction is comprised by interaction of glutamine 493, lysine 417 and glutamine 498 with ACE2 in the wt RBD (Fig3 a,b). Regarding the B.1.1.7 variant glutamine 493 and lysine 417 also contribute majorly to electrostatic interaction between RBD and ACE2. However, glutamine 498, which was displaced from the binding interface, does not contribute to electrostatic interaction as in wt (Fig3 a,b) as markedly reduced electrostatic interaction was calculated (Fig3 c, Tab. 4). Decomposition of electrostatic interaction on the individual residue level for ACE2 reveals that lysine 353 loses its interaction with the RBD in the B.1.1.7 variant (sFig8 a,b, Tab. 5). This fits to our data on contact formation as lysine 353 from ACE2 and glutamine 498 from the RBD lose all their contacts in the B.1.1.7 variant (sFig5 a). In total, we calculated an electrostatic interaction energy of −200 kcal/mol for wt and −165 kcal/mol in B.1.1.7. Computationally more demanding calculations of the binding energy using Molecular Mechanics Generalized Born Surface Area (MM/GBSA) also imply the previously observed reduced electrostatic binding affinity (data not shown). As tyrosine is a bulkier residue than asparagine at position 501, we were now interested in van der Waals contacts also at the level of the individual residues.

**Figure 3:**
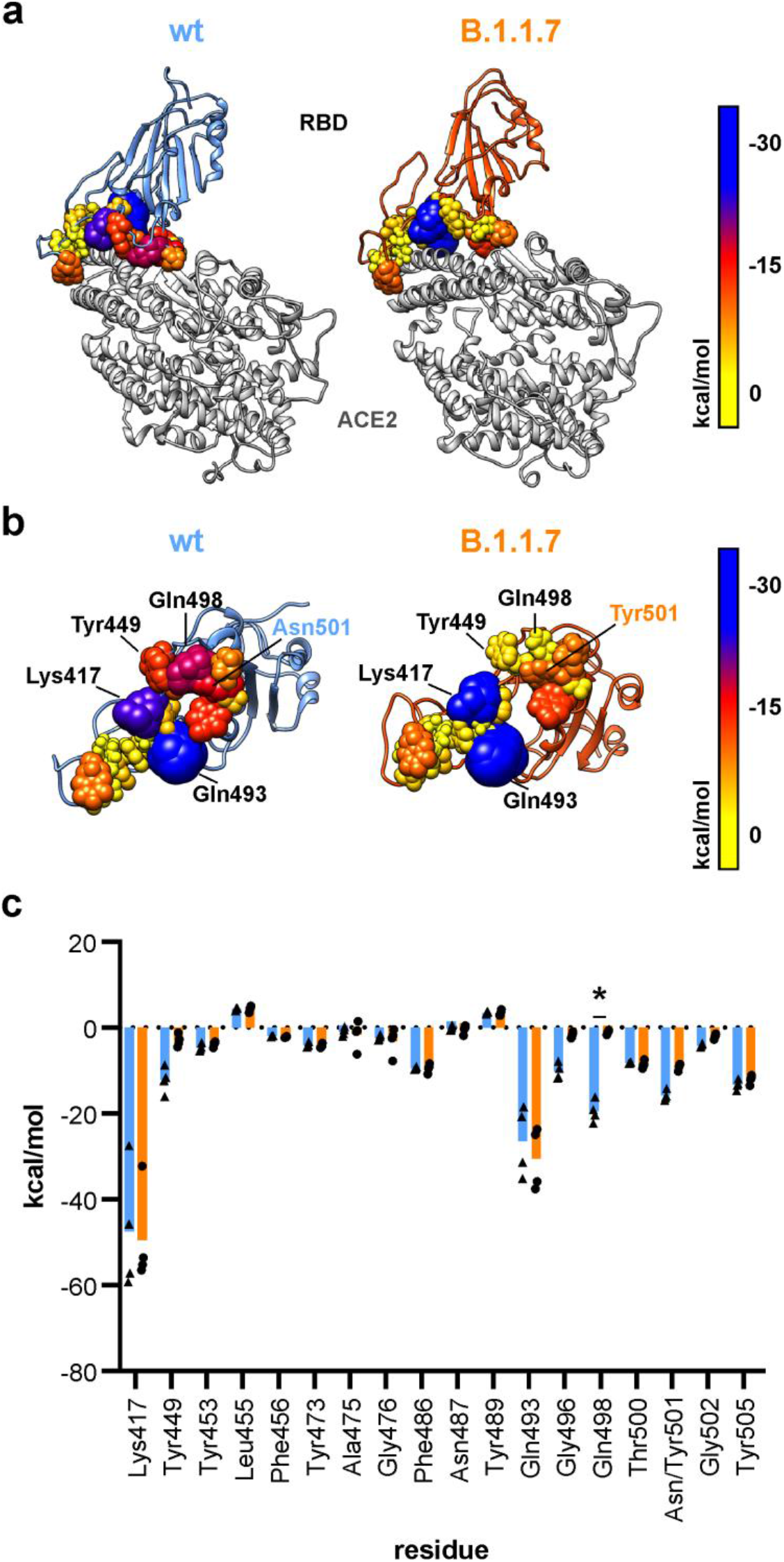
Electrostatic affinity between the RBD and ACE2. a) Structural representation of the RBD (wild type, wt: blue; B.1.1.7: orange) in complex with ACE2 (grey). RBD residues with atoms within a maximum distance of 4 Å from ACE2 are shown as spheres with radii and colors according to their electrostatic linear interaction energy to ACE2. b) View on the interacting interface of the RBD with ACE2. Residue color and atom size represent their electrostatic linear interaction energy to ACE2. c) Quantification of electrostatic linear interaction energy for all residues within 4 Å distance of ACE2 (n = 4; two-way ANOVA; statistical significance assumed for *p<0.05; full list of results in Tab. 4).

**Tab.4:** Statistical analysis of electrostatic interaction energy of residues from the receptor binding domain comparing wt and B.1.1.7 (n = 4; two-way ANOVA; significance assumed for *p<0.05).

| Compare each cell mean with the other cell mean in that row |  |  |  |  |  |
| --- | --- | --- | --- | --- | --- |
| Number of families | 1 |  |  |  |  |
| Number of comparisons per family | 18 |  |  |  |  |
| Alpha | 0,05 |  |  |  |  |
| Sidak's multiple comparisons test | Mean Diff, | 95,00% CI of diff, | Significant? | Summary | Adjusted P Value |
| electrostatic WT - electrostatic UK |  |  |  |  |  |
| Lys417 | 2,002 | -7,629 to 11,63 | No | ns | >0,9999 |
| Tyr449 | -9,196 | -18,83 to 0,4345 | No | ns | 0,0737 |
| Tyr453 | -0,8195 | -10,45 to 8,811 | No | ns | >0,9999 |
| Leu455 | -0,04517 | -9,676 to 9,585 | No | ns | >0,9999 |
| Phe456 | 0,2792 | -9,351 to 9,909 | No | ns | >0,9999 |
| Tyr473 | 0,2544 | -9,376 to 9,885 | No | ns | >0,9999 |
| Ala475 | 0,6749 | -8,955 to 10,31 | No | ns | >0,9999 |
| Gly476 | 0,7864 | -8,844 to 10,42 | No | ns | >0,9999 |
| Phe486 | 0,01221 | -9,618 to 9,643 | No | ns | >0,9999 |
| Asn487 | 0,3207 | -9,310 to 9,951 | No | ns | >0,9999 |
| Tyr489 | 0,06032 | -9,570 to 9,691 | No | ns | >0,9999 |
| Gln493 | 4,088 | -5,543 to 13,72 | No | ns | 0,9775 |
| Gly496 | -8,394 | -18,02 to 1,236 | No | ns | 0,1446 |
| Gln498 | -18,40 | -28,03 to -8,771 | Yes | **** | <0,0001 |
| Thr500 | 0,4097 | -9,221 to 10,04 | No | ns | >0,9999 |
| Asn/Tyr501 | -6,797 | -16,43 to 2,833 | No | ns | 0,4389 |
| Gly502 | -2,077 | -11,71 to 7,553 | No | ns | >0,9999 |
| Tyr505 | -1,129 | -10,76 to 8,502 | No | ns | >0,9999 |

**Tab.5:** Statistical analysis of electrostatic interaction energy of residues from ACE2 comparing wt and B.1.1.7 (n = 4; two-way ANOVA; significance assumed for *p<0.05).

| Compare each cell mean with the other cell mean in that row |  |  |  |  |  |
| --- | --- | --- | --- | --- | --- |
| Number of families | 1 |  |  |  |  |
| Number of comparisons per family | 17 |  |  |  |  |
| Alpha | 0,05 |  |  |  |  |
| Sidak's multiple comparisons test | Mean Diff, | 95,00% CI of diff, | Significant? | Summary | Adjusted P Value |
| electrostatic WT - electrostatic B.1.1.7 |  |  |  |  |  |
| Gln24 | 1,291 | -8,879 to 11,46 | No | ns | >0,9999 |
| Thr27 | 0,08892 | -10,08 to 10,26 | No | ns | >0,9999 |
| Phe28 | 0,09204 | -10,08 to 10,26 | No | ns | >0,9999 |
| Asp30 | 4,740 | -5,430 to 14,91 | No | ns | 0,9401 |
| Lys31 | 0,3279 | -9,841 to 10,50 | No | ns | >0,9999 |
| His34 | -4,305 | -14,47 to 5,864 | No | ns | 0,9740 |
| Glu35 | -1,158 | -11,33 to 9,011 | No | ns | >0,9999 |
| Asp38 | -8,984 | -19,15 to 1,185 | No | ns | 0,1303 |
| Tyr41 | -1,274 | -11,44 to 8,895 | No | ns | >0,9999 |
| Gln42 | -1,510 | -11,68 to 8,659 | No | ns | >0,9999 |
| Met82 | 0,008097 | -10,16 to 10,18 | No | ns | >0,9999 |
| Tyr83 | -0,6152 | -10,78 to 9,554 | No | ns | >0,9999 |
| Asn330 | 0,7600 | -9,409 to 10,93 | No | ns | >0,9999 |
| Lys353 | -28,08 | -38,25 to -17,91 | Yes | **** | <0,0001 |
| Gly354 | -0,5079 | -10,68 to 9,661 | No | ns | >0,9999 |
| Asp355 | -0,8822 | -11,05 to 9,287 | No | ns | >0,9999 |
| Arg357 | 0,2824 | -9,887 to 10,45 | No | ns | >0,9999 |

### Tyrosine at position 501 increases van der Waals interaction locally in B.1.1.7

Decomposition of individual van der Waals linear interaction energies identifies the edges of the RBD-ACE2 interface as important regions. Van der Waals interactions are mainly facilitated by phenylalanine 486, tyrosine 489 and tyrosine 505 in both wt and B.1.1.7 (Fig4 a,b). A major difference was calculated for mutated residue 501 accounting for an asparagine in wt and tyrosine in B.1.1.7 (Fig4 c, Tab. 6). Here we found an increased van der Waals interaction energy in B.1.1.7. Van der Waals interaction energies did not differ in total and were calculated to be around −90 kcal/mol for both wt and B.1.1.7. The van der Waals linear interaction energy was lowered for tyrosine 41 and glutamine 42 from ACE2 (sFig9 a,b, Tab. 7).

**Figure 4:**
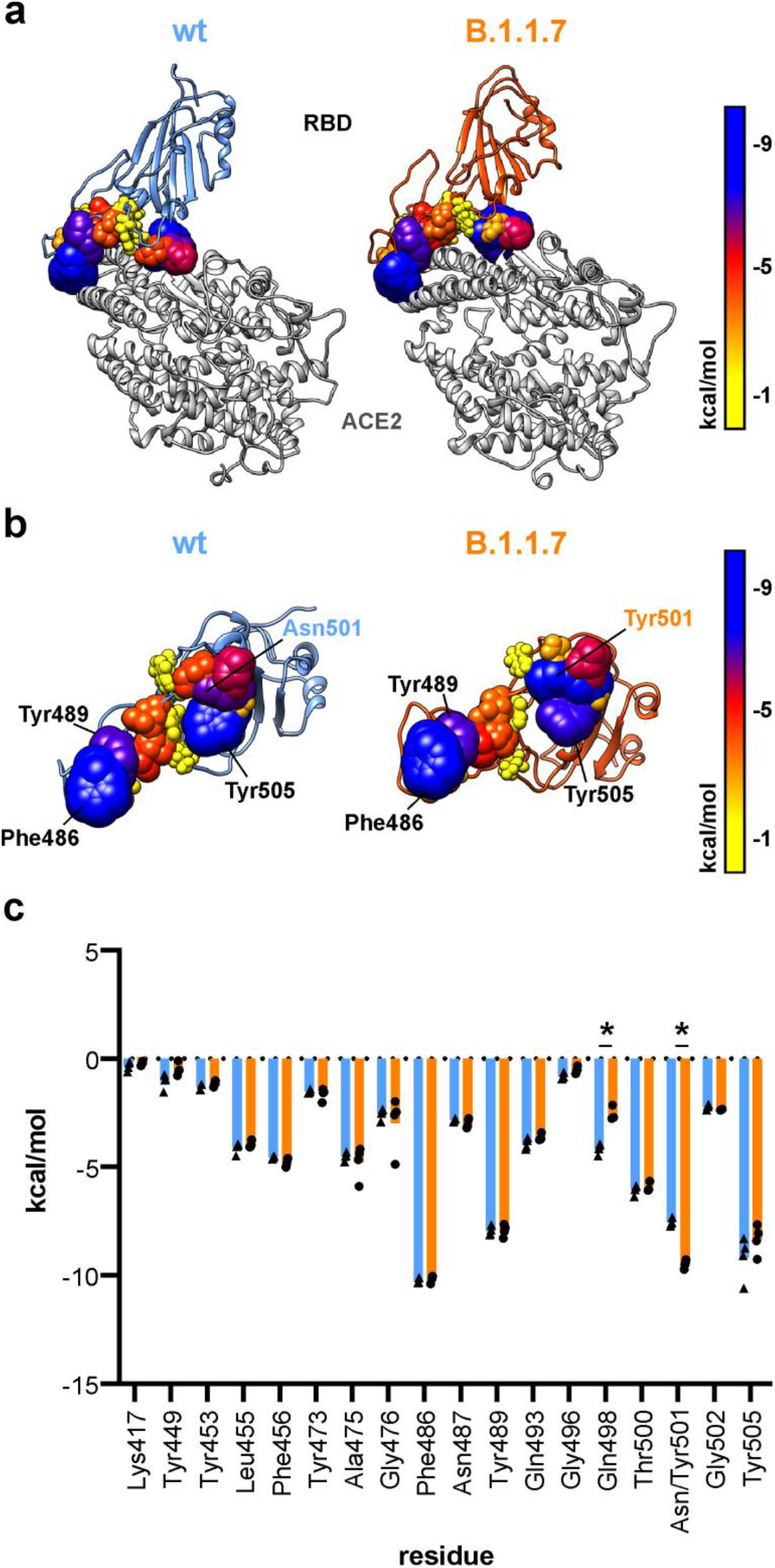
Van der Waals linear interaction energy between the RBD and ACE2. a) Structural representation of the RBD (wild type, wt: blue; B.1.1.7: orange) in complex with ACE2. RBD residues with atoms within a maximum distance of 4 Å from ACE2 are shown as spheres with radii and colors according to their van der Waals linear interaction energy to ACE2. b) View on the interacting interface of the RBD with ACE2. Residue color and atom size represent the van der Waals linear interaction energy to ACE2. c) Quantification of van der Waals linear interaction energy for all RBD residues within 4 Å distance of ACE2 (n = 4; two-way ANOVA; statistical significance assumed for *p<0.05; full list of results in Tab. 6).

**Tab.6:** Statistical analysis of van der Waals interaction energy of residues from the receptor binding domain comparing wt and B.1.1.7 (n = 4; two-way ANOVA; significance assumed for *p<0.05).

| Compare each cell mean with the other cell mean in that row |  |  |  |  |  |
| --- | --- | --- | --- | --- | --- |
| Number of families | 1 |  |  |  |  |
| Number of comparisons per family | 18 |  |  |  |  |
| Alpha | 0,05 |  |  |  |  |
| Sidak's multiple comparisons test | Mean Diff, | 95,00% CI of diff, | Significant? | Summary | Adjusted P Value |
| VdW WT - VdW UK |  |  |  |  |  |
| Lys417 | -0,1159 | -0,8238 to 0,5920 | No | ns | 0,9996 |
| Tyr449 | -0,5646 | -1,674 to 0,5452 | No | ns | 0,5767 |
| Tyr453 | -0,1531 | -0,6207 to 0,3144 | No | ns | 0,9561 |
| Leu455 | -0,1551 | -0,9608 to 0,6506 | No | ns | 0,9995 |
| Phe456 | 0,2132 | -0,4923 to 0,9186 | No | ns | 0,8694 |
| Tyr473 | 0,1292 | -0,9458 to 1,204 | No | ns | >0,9999 |
| Ala475 | 0,2620 | -2,877 to 3,401 | No | ns | >0,9999 |
| Gly476 | 0,4282 | -4,984 to 5,840 | No | ns | >0,9999 |
| Phe486 | -0,04765 | -0,6225 to 0,5272 | No | ns | >0,9999 |
| Asn487 | 0,1392 | -0,6292 to 0,9076 | No | ns | 0,9978 |
| Tyr489 | -0,02050 | -0,9177 to 0,8767 | No | ns | >0,9999 |
| Gln493 | -0,2753 | -1,040 to 0,4894 | No | ns | 0,8832 |
| Gly496 | -0,2742 | -0,7783 to 0,2299 | No | ns | 0,4956 |
| Gln498 | -1,555 | -2,501 to -0,6099 | Yes | ** | 0,0039 |
| Thr500 | -0,1012 | -0,8328 to 0,6305 | No | ns | >0,9999 |
| Asn/Tyr501 | 1,917 | 1,276 to 2,559 | Yes | *** | 0,0001 |
| Gly502 | 0,1328 | -0,2838 to 0,5493 | No | ns | 0,7502 |
| Tyr505 | -0,8373 | -3,986 to 2,311 | No | ns | 0,9887 |

**Tab.7:** Statistical analysis of electrostatic interaction energy of residues from ACE2 comparing wt and B.1.1.7 (n = 4; two-way ANOVA; significance assumed for *p<0.05).

| Compare each cell mean with the other cell mean in that row |  |  |  |  |  |
| --- | --- | --- | --- | --- | --- |
| Number of families | 1 |  |  |  |  |
| Number of comparisons per family | 17 |  |  |  |  |
| Alpha | 0,05 |  |  |  |  |
| Sidak's multiple comparisons test | Mean Diff, | 95,00% CI of diff, | Significant? | Summary | Adjusted P Value |
| electrostatic WT - electrostatic UK |  |  |  |  |  |
| Gln24 | 0,3199 | -2,765 to 3,405 | No | ns | >0,9999 |
| Thr27 | 0,2025 | -0,8693 to 1,274 | No | ns | 0,9924 |
| Phe28 | -0,08288 | -1,123 to 0,9569 | No | ns | >0,9999 |
| Asp30 | -0,1104 | -0,4978 to 0,2770 | No | ns | 0,9745 |
| Lys31 | -0,04280 | -1,378 to 1,292 | No | ns | >0,9999 |
| His34 | -0,5685 | -1,175 to 0,03814 | No | ns | 0,0651 |
| Glu35 | -0,5276 | -1,109 to 0,05356 | No | ns | 0,0772 |
| Asp38 | -0,7532 | -2,592 to 1,086 | No | ns | 0,6362 |
| Tyr41 | -1,162 | -2,035 to -0,2886 | Yes | * | 0,0191 |
| Gln42 | -0,9193 | -1,665 to -0,1739 | Yes | * | 0,0184 |
| Met82 | -0,1952 | -1,267 to 0,8763 | No | ns | 0,9934 |
| Tyr83 | -0,07057 | -0,8227 to 0,6816 | No | ns | >0,9999 |
| Asn330 | 0,1921 | -0,4348 to 0,8190 | No | ns | 0,9457 |
| Lys353 | 0,4709 | -2,538 to 3,480 | No | ns | 0,9951 |
| Gly354 | 0,007700 | -0,6917 to 0,7071 | No | ns | >0,9999 |
| Asp355 | 0,1154 | -0,3833 to 0,6140 | No | ns | 0,9726 |
| Arg357 | 0,1624 | -0,1969 to 0,5216 | No | ns | 0,6094 |

## Discussion

We used molecular dynamics simulations on (i) S protein trimers and (ii) S protein RBD-ACE2 complexes to compare wt and B.1.1.7 variants. Understanding dynamic stability of the S protein is of vital importance to comprehend molecular mechanisms underlying viral entry into the host cell. In particular, this could help to assess superior infectivity of newly emerging viral variants. Although MD simulation data on wt S protein RBD-ACE2 interaction has been reported^19, 20^, data on dynamic changes induced by the N501Y variant found in B.1.1.7 (British), B.1.351 (South African) and P.1 (Brazilian, Japanese) is missing^7^. These three variants also harbor the D614G mutation^7^, which has previously been associated with higher infectivity^5, 21–23^ and viral replication^21, 24^. For variants carrying this D614G mutation it was also suggested that more functional S protein is incorporated into the virion membrane^5^. However, molecular mechanisms explaining the higher infectivity of the D614G mutation are still elusive. Our data derived from simulations of the entire S trimer and comparing wt with B.1.1.7, reveal a rearrangement of salt bridges caused by the loss of interaction between position 614 due to the D614G mutation and lysine 854 in B.1.1.7. This induced flexibility at residues C-terminal of the fusion peptide is enhanced in post-cleavage conformations as also salt bridges formed by aspartate 568 are majorly lost in B.1.1.7. Release of the fusion peptide might allow a faster insertion of the latter into the host cell membrane due to the increased flexibility of the adjacent linker. This might also happen after cleavage at the S2’ position by TMPRSS2^9, 25^ or allow faster entry into host cells after priming the S2’ position by furin during protein secretion^9^, especially as enhanced proteolytic cleavage was proposed for D614G S proteins^26^. Simulation also reveals structural stability of the post-cleavage state induced by salt bridge formation involving arginine 815 that would allow such priming during protein secretion without destabilizing the overall structural integrity. The newly formed salt bridge of aspartate 570 with lysine 964 does not compensate for the loss of aspartate 614 in terms of protein stability, but might rather weaken salt bridges formed by aspartate 568. Additionally, as aspartate 614 comes from the S1 region and lysine 854 from the S2 region of the S protein, dissociation of the S1 and S2 domain of the S protein, important for membrane fusion after entry of the fusion peptide into the host cell membrane, might be enhanced^27^. Furthermore, we show that deletions in the NTD reduce flexibility in the outer region of the S trimer, potentially allowing more spike proteins at the virion host cell interface of B.1.1.7 and thus increasing the probability of viral uptake^5^.

Binding to ACE2 via the receptor-binding domain of the viral S protein is the first step of cellular uptake and infection. Previous reports already highlighted the importance of residues lysine 417, tyrosine 449, glutamine 493 and glutamine 498 on the RBD of the S protein for interface formation with ACE2^7, 20^. At ACE2, residues aspartate 30, lysine 31 and lysine 353 were also identified as important interactors with the RBD of the S protein^7, 20, 28^. We could confirm the importance of the above-mentioned residues for interface formation between RBD and ACE2 here. In addition, our data suggests that the insertion of the bulkier tyrosine at position 501, as found in B.1.1.7, B.1.351 and P.1 excludes glutamine 498 from the RBD-ACE2 interface and alters the molecular architecture of the RBD-ACE2 interface of the S protein. We measured a reduced electrostatic affinity of glutamine 498 from the S protein to residues on ACE2. For tyrosine 501 from the S protein RBD we found an increased van der Waals interaction with lysine 353 (ACE2) in the N501Y variant. Similar RMSD values of the RBD for wt and B.1.1.7 suggest that amino acid deletions/exchanges in the NTD and fusion peptide adjacent loop region cause local changes and do not change the overall flexibility of the RBD. In conclusion, our data suggest a higher mobility of the fusion peptide after release and reduced electrostatic affinity of the S protein RBD to ACE2 in B.1.1.7. This combination could allow faster membrane fusion of the virion with the host cell membrane and could be explained by (i) faster insertion of the fusion peptide and (ii) faster dissociation of the S protein RBD-ACE2 complex due to reduced electrostatic affinity. Dissociation of the S protein components is important for efficient membrane fusion after insertion of the fusion peptide. Our data might serve as a basis for advanced biochemical and cell biological experiments to better understand the importance of individual amino acid exchanges for viral pathology.

## Methods

### Generation of the starting structures

To generate starting structures for wild type, wild type with the D614G mutation (wt+D614G) and B.1.1.7 SARS-CoV-2 we used the protein sequence annotated in UniProt (www.uniprot.org) with the identifier P0DTC2 (SARS-CoV-2; wild-type) and changed the sequence in accordance with the reported deletions (del69-70, del144) for the SARS-CoV-2 B.1.1.7 variant. Using Swiss-Model Expasy (www.swissmodel.expasy.org) we generated a model for the SARS-CoV-2 wild type, wild type with the D614G mutation and SARS-CoV-2 B.1.1.7 variant on the basis of the Cryo-EM solved structure of the triple ACE2 bound SARS-CoV-2 spike protein trimer (PDB ID: 7KMS)^16^ and used chain A as a template. After modeling, structures were transferred into UCSF Chimera^29^ and single amino acid exchanges were inserted into the B.1.1.7 structure using the *swapaa* command. These were N501Y, A570D, D614G, P681H, T716I, S982A and D1118H. Post-translational modifications (here sugars) were removed. Monomeric spike proteins of wt, wt+D614G and B.1.1.7 were then C3 symmetrized to generate trimeric spike protein complexes and used as starting structures for molecular dynamics (MD) simulations. Cleavage of wild type and the B.1.1.7 variant was structurally implemented in the PDB files used as starting structures by adding TER records (indicates end of a chain) between arginine 815 and serine 816 in all three chains of the trimer.

To investigate the interface between the RBD of the spike protein and ACE2, the respective wild type starting structure was obtained from the PDB database (PDB ID code: 7KMB)^16^. In order to generate also the starting structure for the MD simulations of the B.1.1.7 variant the N501Y amino acid exchange was introduced with Swiss-PdbViewer 4.1.0.

### Molecular dynamics simulations

Molecular dynamics simulations were performed using version 20 of the Amber Molecular Dynamics software package (ambermd.org)^30^ and the ff14SB force field^31^. With the Amber Tool LEaP, all systems were electrically neutralized with Na^+^ ions and solvated with TIP3P^32^ water molecules. The trimeric SARS CoV-2 spike protein was solvated in a cuboid water box with at least 15 Å distance from the borders to the solute, whereas the receptor-binding domain (RBD) complexed with ACE2 was solvated in a water box with the shape of a truncated octahedron and at least 25 Å distance from the borders to the solute.

The simulations followed a previously applied protocol^33^. At first, a minimization was carried out in three subsequent steps to optimize the geometry of the starting structures. In the first step of the minimization, the water molecules were minimized, while all remaining atoms were restrained with a constant force of 10 kcal·mol^−1^·Å^−2^ to the initial positions. In the second step, additional relaxation of the sodium ions and the hydrogen atoms of the protein were allowed, while the remaining protein was restrained with 10 kcal·mol^−1^·Å^−2^. In the last step, no restraints were used, so that the whole protein, the ions, and the water molecules were minimized. All three minimization parts started with 2500 steps using the steepest descent algorithm, followed by 2500 steps of a conjugate gradient minimization. After the minimization, the systems were equilibrated in two successive steps. In the first step, the temperature was raised from 10 to 310 K within 0.1 ns and the protein was restrained with a constant force of 5 kcal·mol^−1^·Å^−2^. In the second step (0.4 ns length), only the Cα atoms of the protein were restrained with a constant force of 5 kcal·mol^−1^·Å^−2^. In both equilibration steps, the time step was 2 fs. Minimization and equilibration were carried out on CPUs, whereas the subsequent production runs were performed using pmemd.CUDA on Nvidia A100 GPUs^34–36^. In the following, 200 ns (trimeric spike protein) or 500 ns (RBD complexed with ACE2) long production runs were conducted without any restraints and at 310 K (regulated by a Berendsen thermostat^37^). Furthermore, the constant pressure periodic boundary conditions were used with an average pressure of 1 bar and isotropic position scaling. For bonds involving hydrogen, the SHAKE algorithm^38^ was applied in the equilibration and production phase. In order to accelerate the production phase of the molecular dynamics (MD) simulations, hydrogen mass repartitioning (HMR)^39^ was used in combination with a time step of 4 fs. The MD simulations were performed two times for all forms of the trimeric spike protein and four times for both forms of the RBD in complex with ACE2.

Trajectory analysis (analysis of root-mean-square deviation of atomic positions (RMSD), root-mean-square fluctuations (RMSF), analysis of native contacts, measurement of interatomic distances, calculation of linear interaction energy (electrostatic and van der Waals interactions, used default cutoff of 12.0 Å) was carried out using the Amber tool cpptraj^40^. Contacts were evaluated with an in-house Perl script parsing the trajectory using the prior named Amber tools and assigning contacts based on a distance criterion of ≤5 Å between any pair of atoms, as it was done previously^41^.

### Statistics and display

Statistical analyses were generated with GraphPad Prism (version 8.0.0 for Windows, GraphPad Software, San Diego, California USA, www.graphpad.com) and statistical tests were applied as indicated below the figure. Plots were generated in GraphPad and Gnuplot (version 4.6). All structure images were made with UCSF Chimera 1.15^29^.

## Acknowledgements

The authors gratefully acknowledge the compute resources and support provided by the Erlangen Regional Computing Center (RRZE) and by NHR@FAU.

## Author contributions

E.S., F.Z., P.A. conceived the study; E.S. conducted the MD simulations; E.S., M.C., P.A. performed data analysis; E.S., M.C., L.H., P.A. generated visualization of the data; E.S., H.S., F.P., F.Z., P.A. contributed to the design of the study as well as discussion of the data; E. S., F.Z., P.A. wrote the initial draft; all authors reviewed the manuscript prior to submission.

---

**sFig1:**
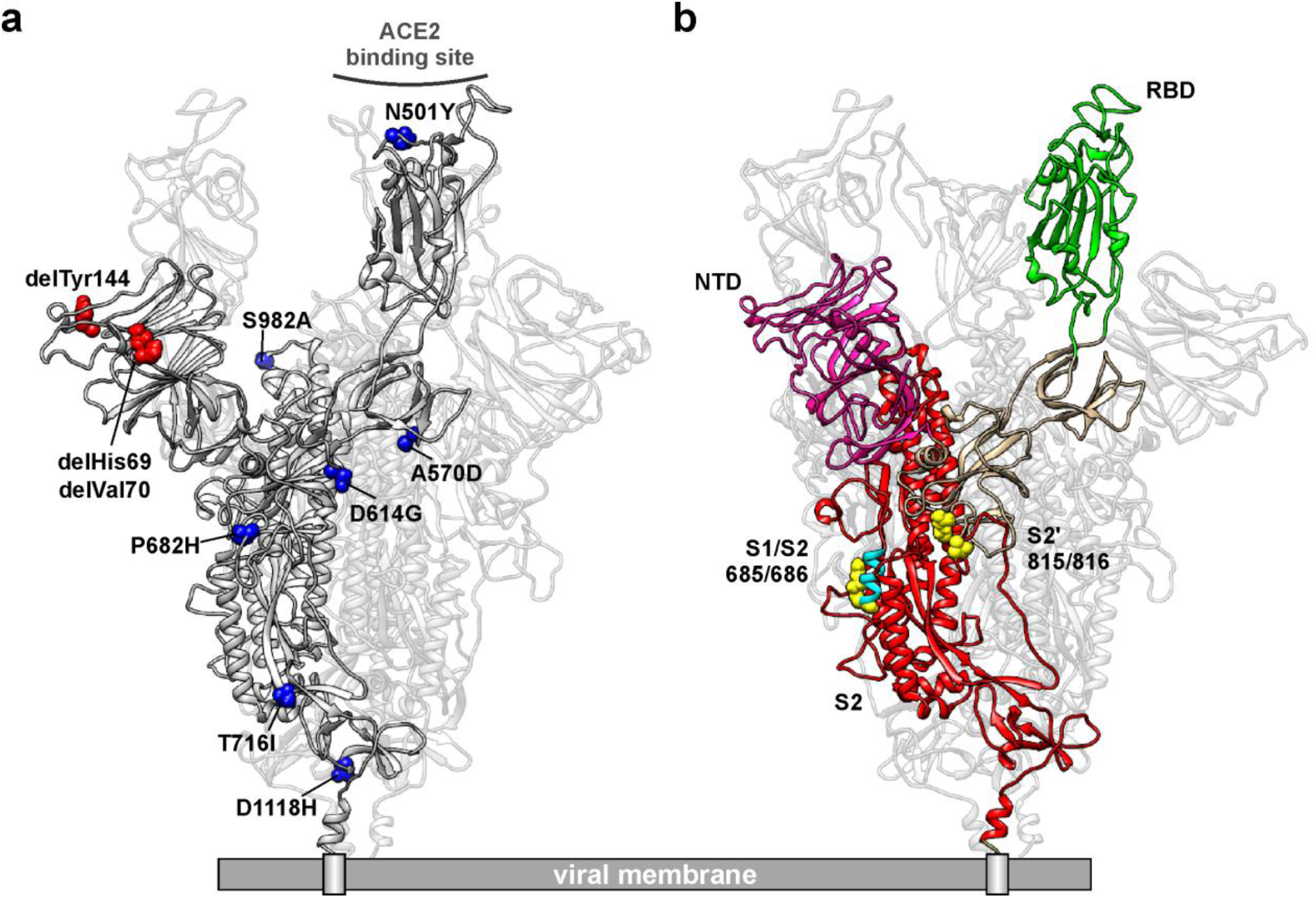
a) Structural representation of the SARS-CoV-2 S protein on the viral surface with amino acid residues shown as spheres that are deleted (red) or mutated (blue) in the B.1.1.7 variant. b) Structural representation of the SARS-CoV-2 S protein on the viral surface with domains highlighted in color as in figure 1a (N-terminal domain, NTD in pink; receptor-binding domain, RBD in green; S1 region in khaki; S1/S2 and S2’ cleavage sites in yellow with amino acids as spheres; fusion peptide in cyan; S2 domain in red).

**sFig2:**
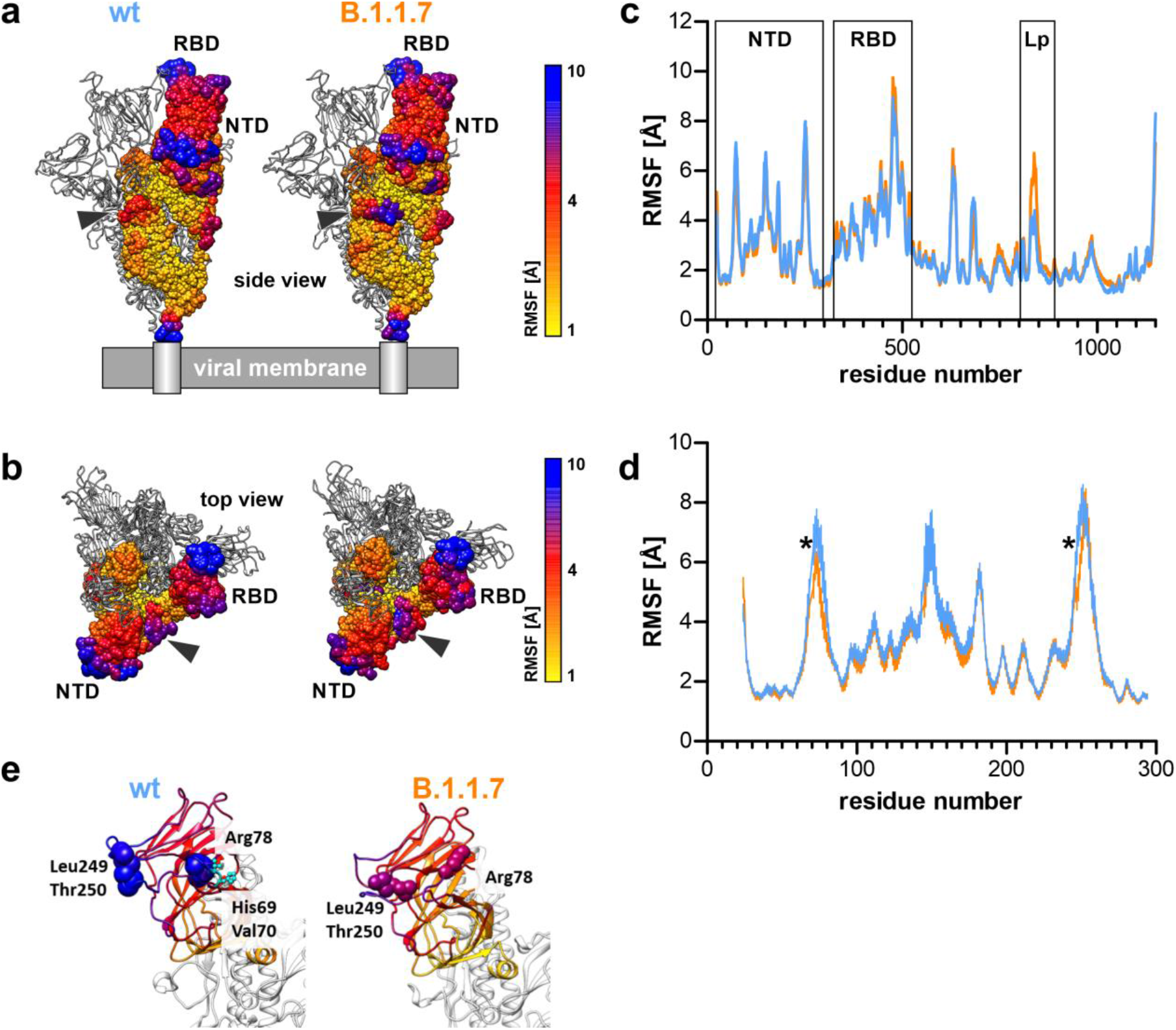
a) Structural representation of the SARS-CoV-2 S protein in side view on the viral surface with residues shown as colored spheres for one subunit according to their RMSF value (grey arrowhead, flexible loop region in B.1.1.7). b) Top view of a). c) Line plot of RMSF values for calculated averages of six subunits of the wt (blue) and B.1.1.7 (orange) S protein. d) Line plot of RMSF values averaged over six subunits of the N-terminal domain from wt (blue) and B.1.1.7 (orange). Asterisks indicate positions of significant differences (n = 6; two-way ANOVA; significance assumed for p<0.05; full list in Tab. 1). e) Structural representation of the N-terminal domain colored according to RMSF values. Residues that differ significantly in RMSF values are represented as spheres. Additionally, His69 and Val70 are shown for wild type (wt) and are missing as del69,70 in B.1.1.7 (n = 6; two-way ANOVA; significance assumed for *p<0.05; full list in Tab. 1). NTD = N-terminal domain, RBD = receptor-binding domain, Lp = loop region around amino acids 836-844.

**sFig3:**
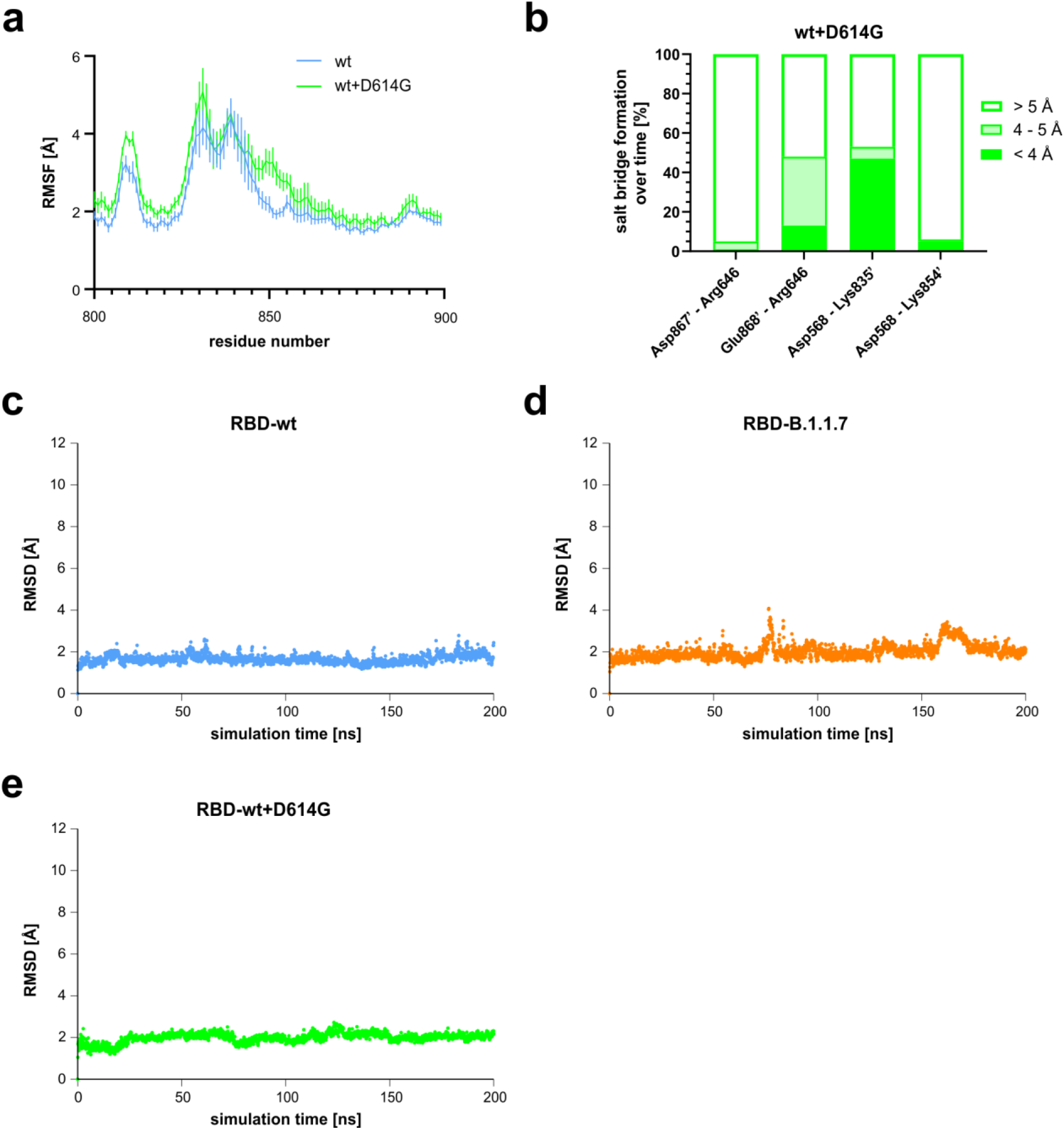
a) Line plot of root-mean-square fluctuation (RMSF) values for amino acid residues 800 to 900. Averages were calculated for all three trimer subunits and both simulation runs of the wild type (wt, blue) and wt+D614G variant (green). b) Percentage of salt bridge formation over time for four different residue pairs. All residue pairs represent interchain interactions with residues from two different, but directly neighboring chains within the trimeric spike protein. c-e) Conformational stability was measured as root-mean-square deviation (RMSD) values over simulation time for the receptor-binding domain (RBD) of c) the wild type d) the B.1.1.7 variant and e) the wild type with the D614G mutation. One representative plot is shown for wild type spike protein and its variants.

**sFig4:**
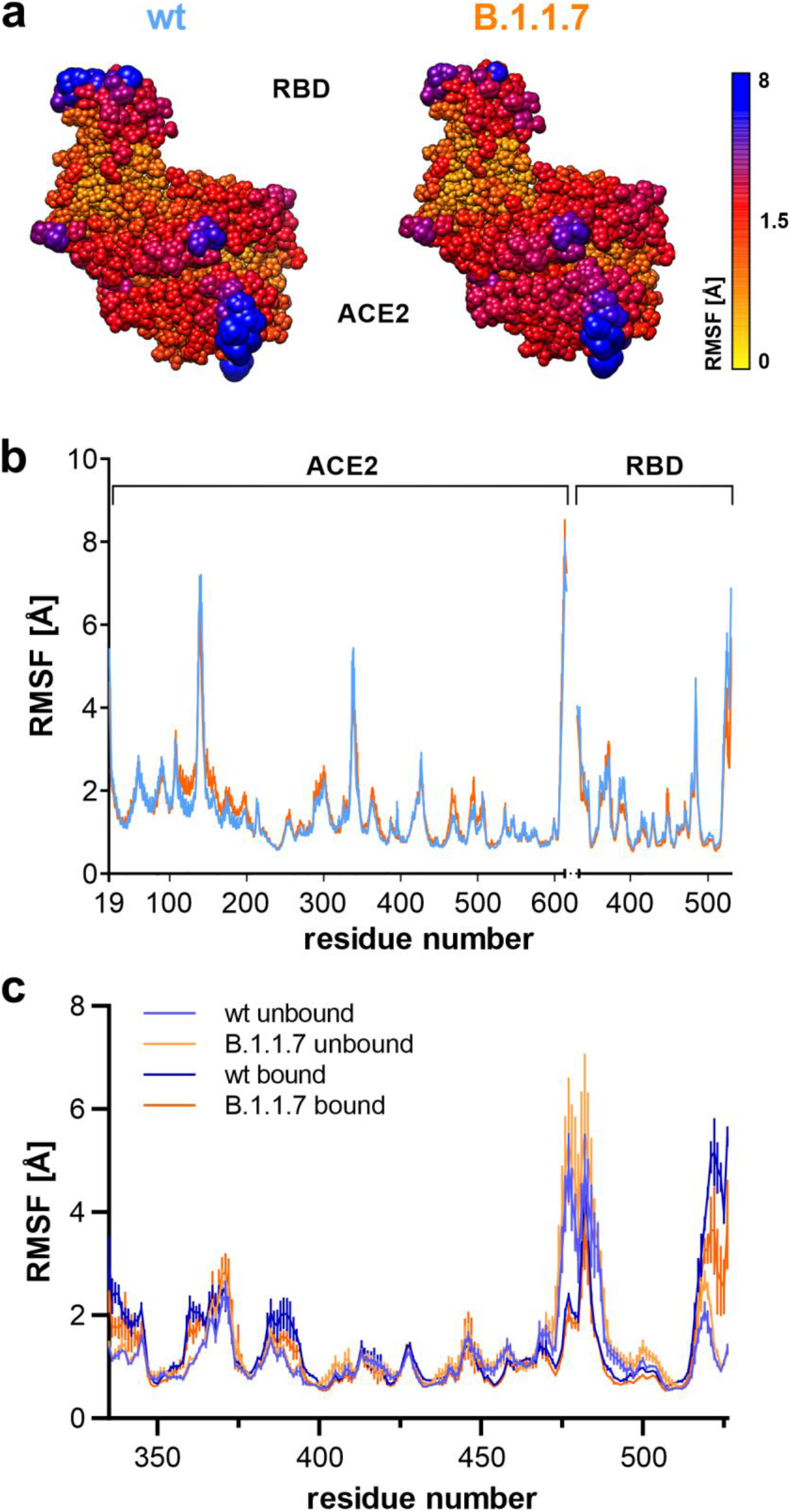
a) Structural flexibility of the receptor-binding domain-ACE2 (RBD-ACE2) complex. Residues are colored and shown as spheres of different size according to their individual RMSF values. RBD denotes the receptor-binding domain and ACE2 the angiotensin-converting enzyme 2. b) Line plot of RMSF values averaged over four molecular dynamics (MD) simulation runs for the RBD-ACE2 complex with wild type (wt) in blue and the B.1.1.7 variant in orange. c) Line plot of RMSF values from the RBD in the unbound state (from the S protein trimer MD simulations) and in the bound state to ACE2 (from the RBD-ACE2 complex MD simulations).

**sFig5:**
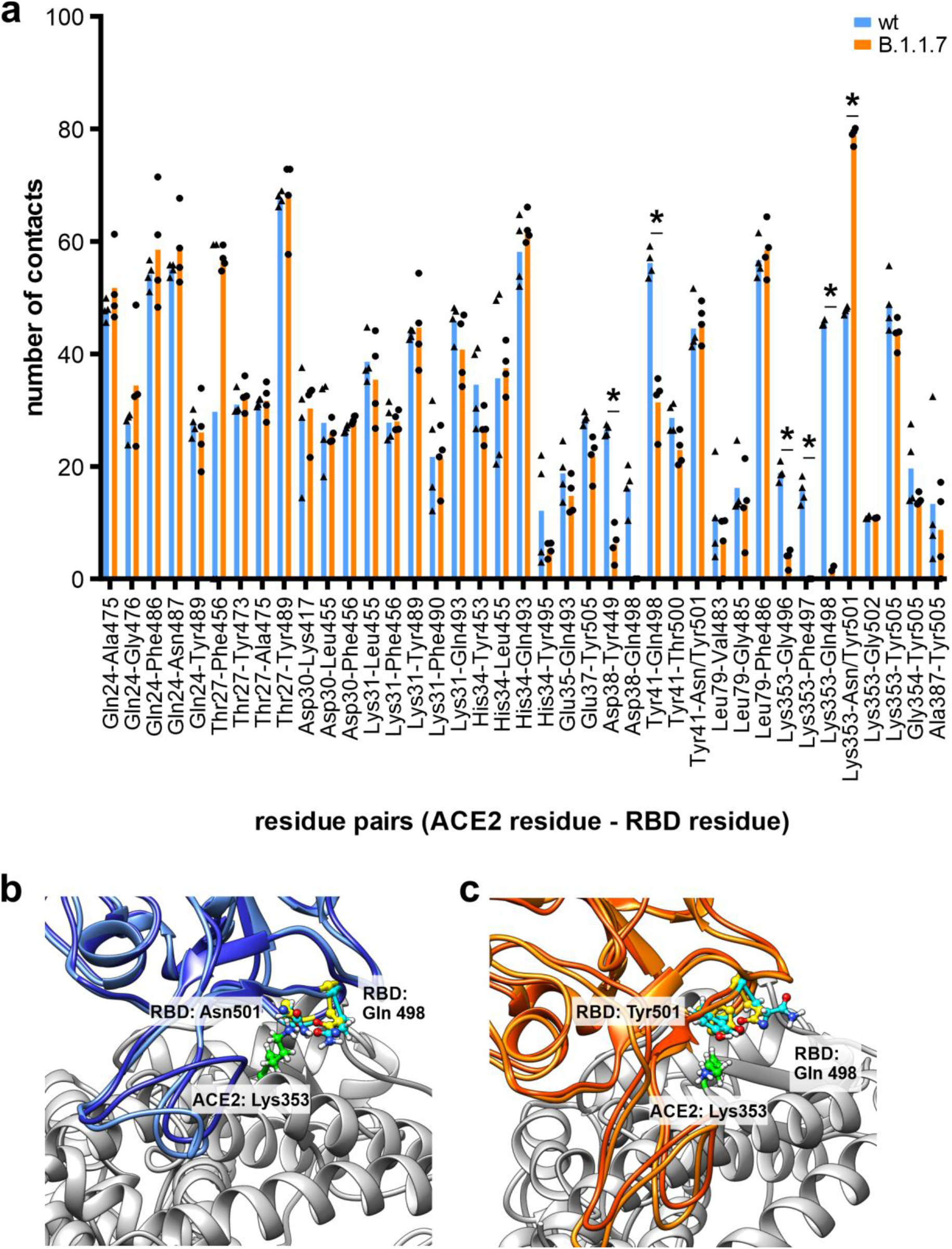
Average number of contacts for residues from ACE2 and the receptor-binding domain (RBD). Residues were included when they came in closer proximity than 5 Å. Loss of contacts were calculated for glutamine 498 (Gln498) in B.1.1.7, and gain of contacts for the newly inserted tyrosine at position 501 (Tyr501) with lysine 353 (Lys353). Wild type (wt) is shown in blue and B.1.1.7 in orange (n = 4, two-way ANOVA; statistical significance was assumed for *p<0.05; full list of results in Tab. 3). b) Structure of the RBD-ACE2 interface with residues asparagine 501 (Asn501; wild type in blue) or tyrosine 501 (Tyr501; B.1.1.7 in orange), glutamine 498 (Gln498) (all from RBD) and lysine 353 (green) from ACE2 shown in ball-and-stick. Yellow residues represent the starting conformation and residues in cyan are representative residue side chain conformations acquired during molecular dynamics simulation.

**sFig6:**
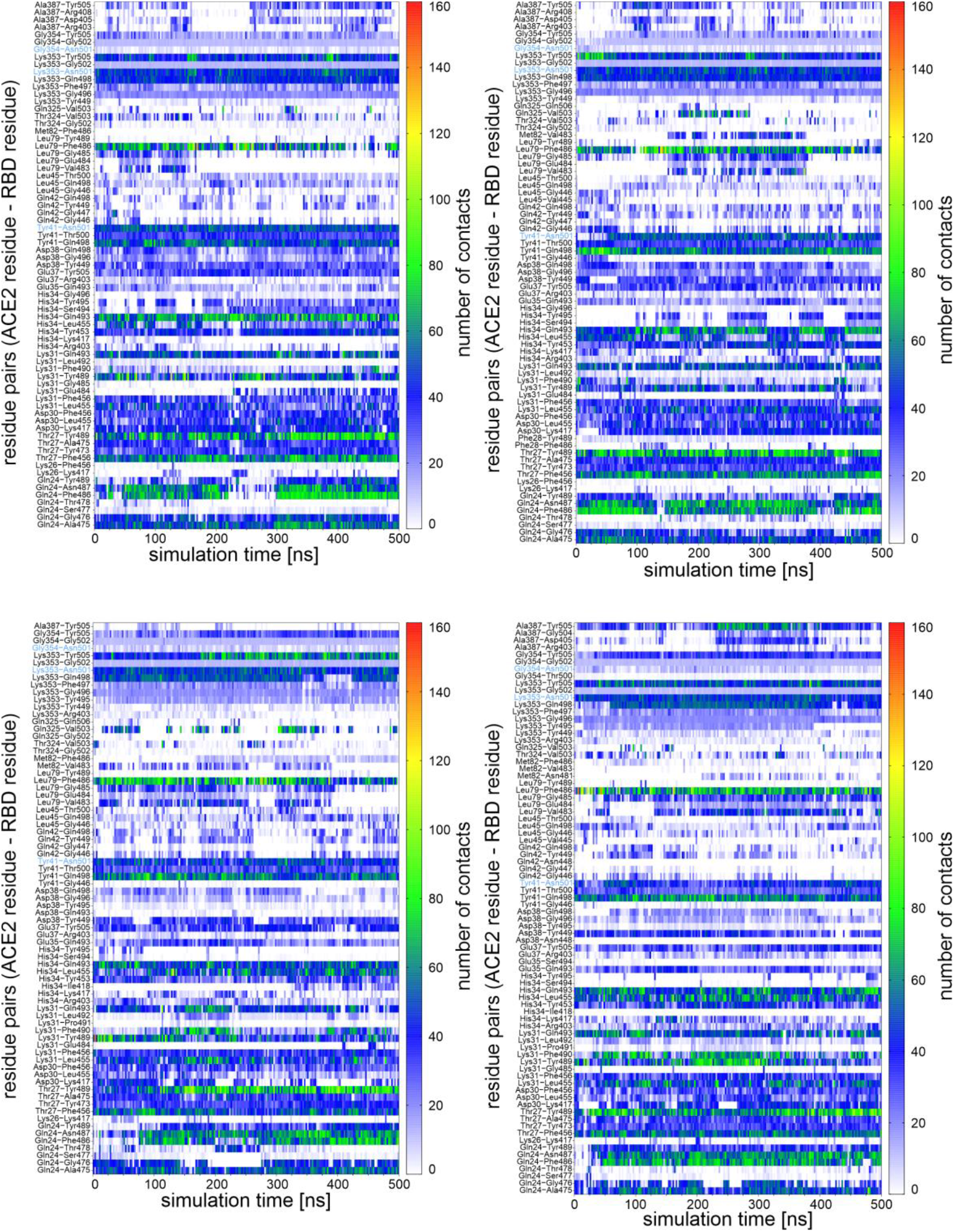
Individual intermolecular contact plots for all four molecular dynamics (MD) simulation runs of the wild type RBD-ACE2 complex. The number of contacts, calculated as the number of interresidue atom pairs that are within a maximum distance of 5 Å from each other, was plotted color-coded for intermolecular residue pairs over the simulation time. For instance, the residue pair Lys353-Asn501, with Lys353 expressed on ACE2 and Asn501 expressed on the RBD, has in all four MD simulations a calculated number of contacts between 40 and 60 over the whole simulation time. Residue pairs with asparagine 501 are highlighted in blue.

**sFig7:**
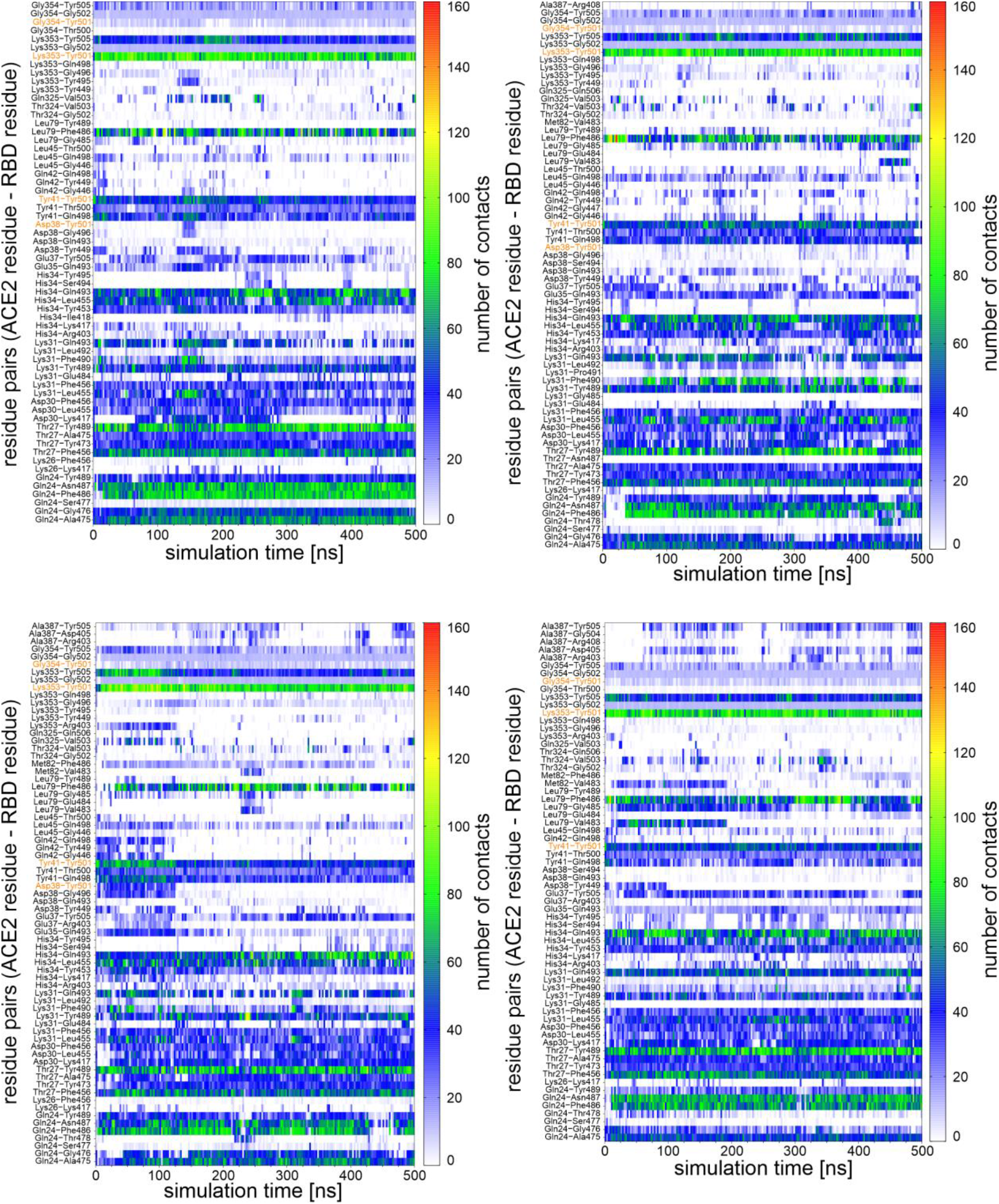
Individual intermolecular contact plots for all four molecular dynamics (MD) simulation runs of the B.1.1.7 RBD-ACE2 complex. The number of contacts, calculated as the number of interresidue atom pairs that are within a maximum distance of 5 Å from each other, was plotted color-coded for intermolecular residue pairs over the simulation time. For instance, the residue pair Lys353-Tyr501, with Lys353 expressed on ACE2 and Tyr501 expressed on the mutated RBD, has in all four MD simulations a calculated number of contacts around 80 over the whole simulation time. Residue pairs with mutated tyrosine 501 are highlighted in orange.

**sFig8:**
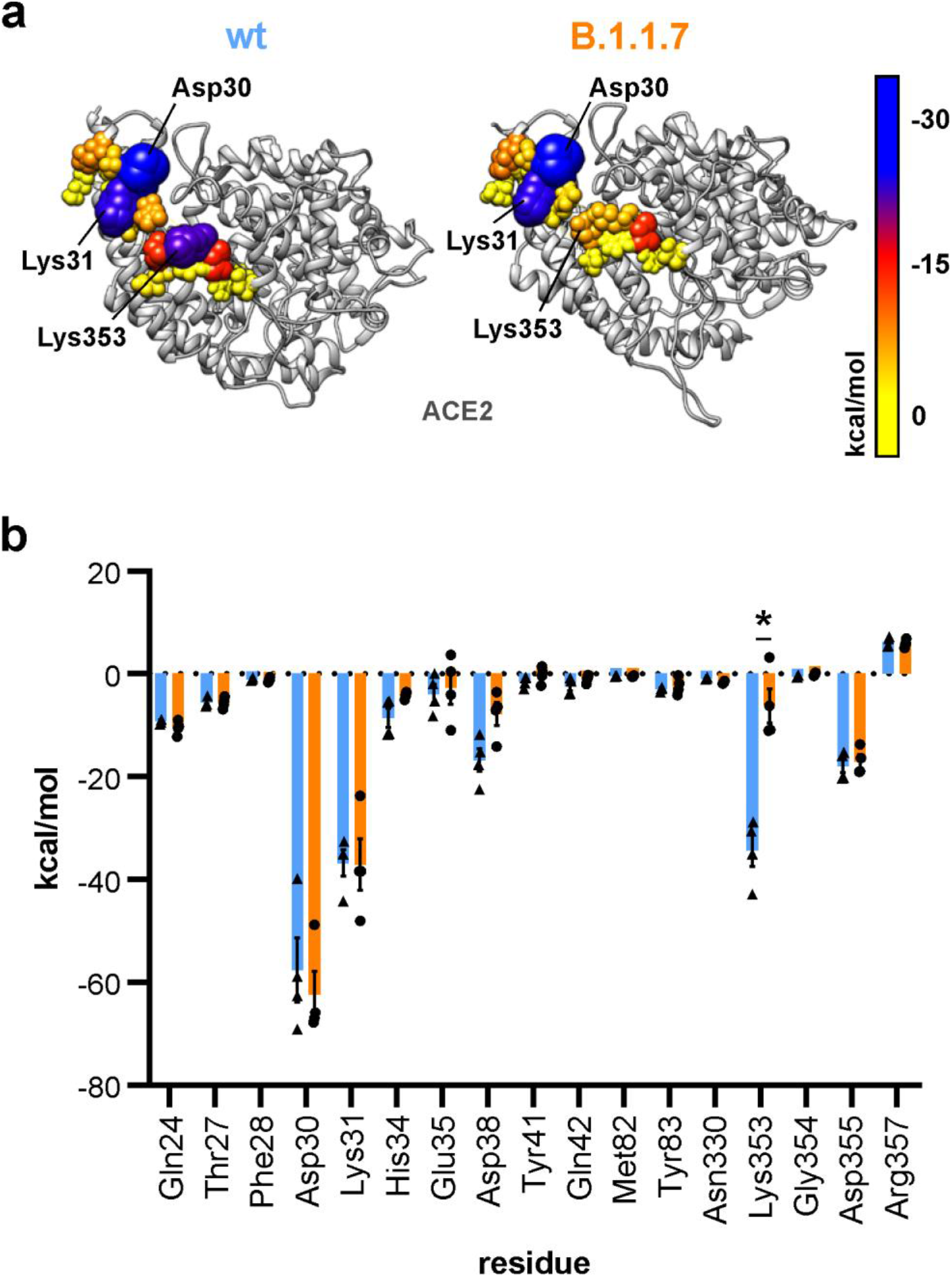
a) Electrostatic affinity shown on the RBD-ACE2 interface for ACE2 residues. Residues in closer contact than 4 Å are shown as spheres of different size and color according to their electrostatic affinity with the RBD. b) Quantification of the electrostatic linear interaction energy for all residues within a radius of 4 Å around the receptor-binding domain (n =4; two-way ANOVA; statistical significance was assumed for *p<0.05; full list of results in Tab. 5).

**sFig9:**
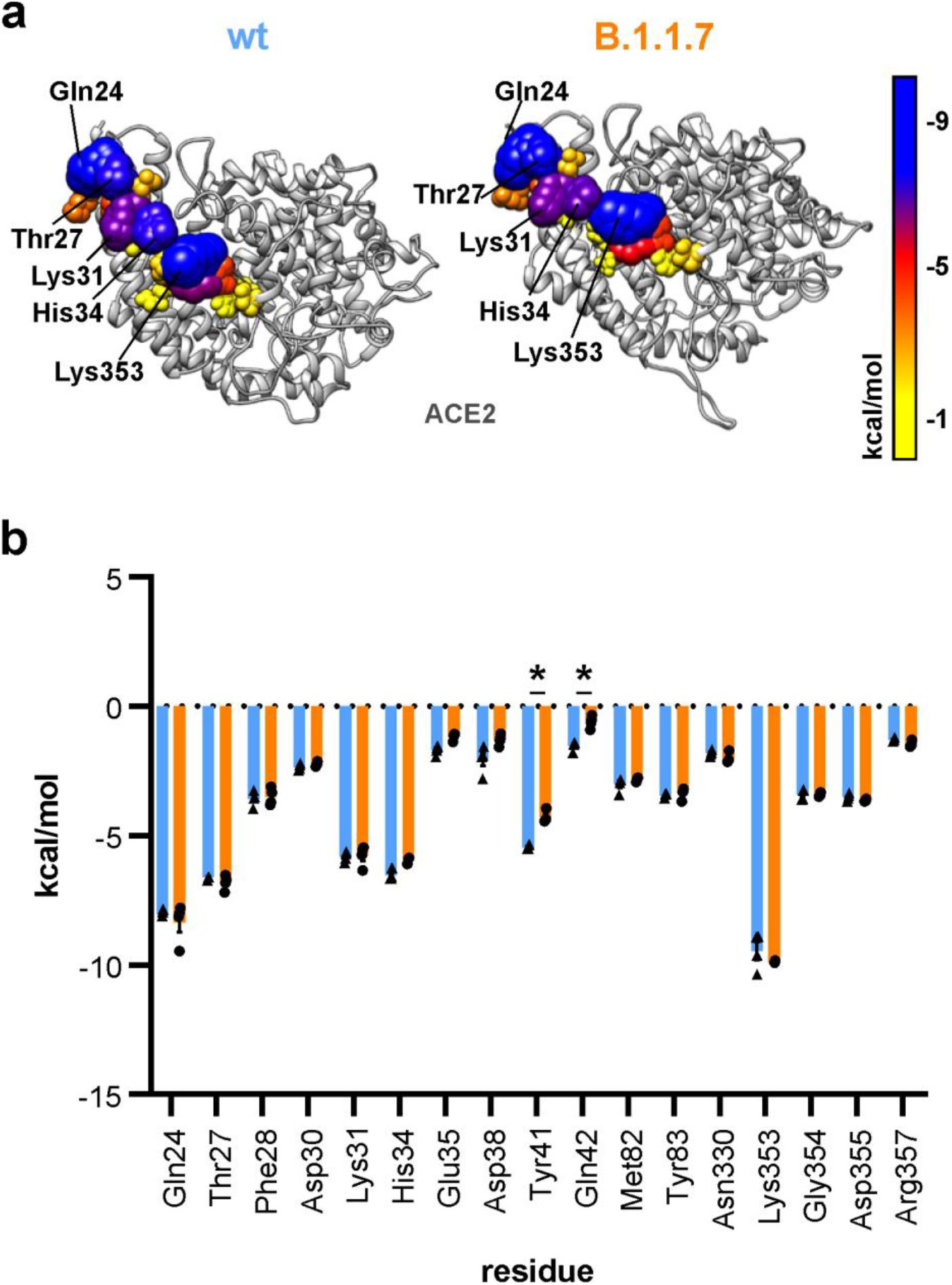
a) Van der Waals linear interaction energy shown on the RBD-ACE2 interface for ACE2 residues. Residues in closer contact than 4 Å are shown as spheres of different size and color according to their van der Waals linear interaction energy with the RBD. b) Quantification of the van der Waals linear interaction energy for all residues within a radius of 4 Å around the receptor-binding domain (n =4; two-way ANOVA; statistical significance was assumed for *p<0.05; full list of results in Tab. 7).

## Literature

1. https://covid19.who.int/. World Healt Organization Coronavirus Dashboard

2. Galloway, S. E.; Paul, P.; MacCannell, D. R.; Johansson, M. A.; Brooks, J. T.; MacNeil, A.; Slayton, R. B.; Tong, S.; Silk, B. J.; Armstrong, G. L.; Biggerstaff, M.; Dugan, V. G., Emergence of SARS-CoV-2 B.1.1.7 Lineage - United States, December 29, 2020-January 12, 2021. MMWR Morb Mortal Wkly Rep 2021, 70 (3), 95–99.

3. Leung, K.; Shum, M. H.; Leung, G. M.; Lam, T. T.; Wu, J. T., Early transmissibility assessment of the N501Y mutant strains of SARS-CoV-2 in the United Kingdom, October to November 2020. Euro Surveill 2021, 26 (1).

4. Korber, B.; Fischer, W. M.; Gnanakaran, S.; Yoon, H.; Theiler, J.; Abfalterer, W.; Hengartner, N.; Giorgi, E. E.; Bhattacharya, T.; Foley, B.; Hastie, K. M.; Parker, M. D.; Partridge, D. G.; Evans, C. M.; Freeman, T. M.; de Silva, T. I.; Sheffield, C.-G. G.; McDanal, C.; Perez, L. G.; Tang, H.; Moon-Walker, A.; Whelan, S. P.; LaBranche, C. C.; Saphire, E. O.; Montefiori, D. C., Tracking Changes in SARS-CoV-2 Spike: Evidence that D614G Increases Infectivity of the COVID-19 Virus. Cell 2020, 182 (4), 812–827 e19.

5. Zhang, L.; Jackson, C. B.; Mou, H.; Ojha, A.; Peng, H.; Quinlan, B. D.; Rangarajan, E. S.; Pan, A.; Vanderheiden, A.; Suthar, M. S.; Li, W.; Izard, T.; Rader, C.; Farzan, M.; Choe, H., SARS-CoV-2 spike-protein D614G mutation increases virion spike density and infectivity. Nat Commun 2020, 11 (1), 6013.

6. Nicholas G. Davies, R. C. B., Christopher I. Jarvis, Adam J. Kucharski, James Munday, Carl A. B. Pearson, Timothy W. Russell, Damien C. Tully, Sam Abbott, Amy Gimma, William Waites, Kerry LM Wong, Kevin van Zandvoort, CMMID COVID-19 Working Group, Rosalind M. Eggo, Sebastian Funk, Mark Jit, Katherine E. Atkins, W. John Edmunds, Estimated transmissibility and severity of novel SARS-CoV-2 Variant of Concern 202012/01 in England. medRxiv 2020.12.24.20248822; doi: https://doi.org/10.1101/2020.12.24.20248822 2020.

7. Li, F.; Li, W.; Farzan, M.; Harrison, S. C., Structure of SARS coronavirus spike receptor-binding domain complexed with receptor. Science 2005, 309 (5742), 1864–8.

8. Zhou, P.; Yang, X. L.; Wang, X. G.; Hu, B.; Zhang, L.; Zhang, W.; Si, H. R.; Zhu, Y.; Li, B.; Huang, C. L.; Chen, H. D.; Chen, J.; Luo, Y.; Guo, H.; Jiang, R. D.; Liu, M. Q.; Chen, Y.; Shen, X. R.; Wang, X.; Zheng, X. S.; Zhao, K.; Chen, Q. J.; Deng, F.; Liu, L. L.; Yan, B.; Zhan, F. X.; Wang, Y. Y.; Xiao, G. F.; Shi, Z. L., A pneumonia outbreak associated with a new coronavirus of probable bat origin. Nature 2020, 579 (7798), 270–273.

9. Hoffmann, M.; Kleine-Weber, H.; Schroeder, S.; Kruger, N.; Herrler, T.; Erichsen, S.; Schiergens, T. S.; Herrler, G.; Wu, N. H.; Nitsche, A.; Muller, M. A.; Drosten, C.; Pohlmann, S., SARS-CoV-2 Cell Entry Depends on ACE2 and TMPRSS2 and Is Blocked by a Clinically Proven Protease Inhibitor. Cell 2020, 181 (2), 271–280 e8.

10. Shang, J.; Wan, Y.; Luo, C.; Ye, G.; Geng, Q.; Auerbach, A.; Li, F., Cell entry mechanisms of SARS-CoV-2. Proc Natl Acad Sci U S A 2020, 117 (21), 11727–11734.

11. Belouzard, S.; Chu, V. C.; Whittaker, G. R., Activation of the SARS coronavirus spike protein via sequential proteolytic cleavage at two distinct sites. Proc Natl Acad Sci U S A 2009, 106 (14), 5871–6.

12. Ou, X.; Zheng, W.; Shan, Y.; Mu, Z.; Dominguez, S. R.; Holmes, K. V.; Qian, Z., Identification of the Fusion Peptide-Containing Region in Betacoronavirus Spike Glycoproteins. J Virol 2016, 90 (12), 5586–5600.

13. Hoffmann, M.; Kleine-Weber, H.; Pohlmann, S., A Multibasic Cleavage Site in the Spike Protein of SARS-CoV-2 Is Essential for Infection of Human Lung Cells. Mol Cell 2020, 78 (4), 779–784 e5.

14. Wrapp, D.; Wang, N.; Corbett, K. S.; Goldsmith, J. A.; Hsieh, C. L.; Abiona, O.; Graham, B. S.; McLellan, J. S., Cryo-EM Structure of the 2019-nCoV Spike in the Prefusion Conformation. bioRxiv 2020.

15. Walls, A. C.; Park, Y. J.; Tortorici, M. A.; Wall, A.; McGuire, A. T.; Veesler, D., Structure, Function, and Antigenicity of the SARS-CoV-2 Spike Glycoprotein. Cell 2020, 183 (6), 1735.

16. Zhou, T.; Tsybovsky, Y.; Gorman, J.; Rapp, M.; Cerutti, G.; Chuang, G. Y.; Katsamba, P. S.; Sampson, J. M.; Schon, A.; Bimela, J.; Boyington, J. C.; Nazzari, A.; Olia, A. S.; Shi, W.; Sastry, M.; Stephens, T.; Stuckey, J.; Teng, I. T.; Wang, P.; Wang, S.; Zhang, B.; Friesner, R. A.; Ho, D. D.; Mascola, J. R.; Shapiro, L.; Kwong, P. D., Cryo-EM Structures of SARS-CoV-2 Spike without and with ACE2 Reveal a pH-Dependent Switch to Mediate Endosomal Positioning of Receptor-Binding Domains. Cell Host Microbe 2020, 28 (6), 867–879 e5.

17. Lan, J.; Ge, J.; Yu, J.; Shan, S.; Zhou, H.; Fan, S.; Zhang, Q.; Shi, X.; Wang, Q.; Zhang, L.; Wang, X., Structure of the SARS-CoV-2 spike receptor-binding domain bound to the ACE2 receptor. Nature 2020, 581 (7807), 215–220.

18. CDC https://www.cdc.gov/coronavirus/2019-ncov/cases-updates/variant-surveillance/variant-info.html#Concern. (accessed 2021-03-17).

19. Yang, J.; Petitjean, S. J. L.; Koehler, M.; Zhang, Q.; Dumitru, A. C.; Chen, W.; Derclaye, S.; Vincent, S. P.; Soumillion, P.; Alsteens, D., Molecular interaction and inhibition of SARS-CoV-2 binding to the ACE2 receptor. Nat Commun 2020, 11 (1), 4541.

20. Ali, A.; Vijayan, R., Dynamics of the ACE2-SARS-CoV-2/SARS-CoV spike protein interface reveal unique mechanisms. Sci Rep 2020, 10 (1), 14214.

21. Hou, Y. J.; Chiba, S.; Halfmann, P.; Ehre, C.; Kuroda, M.; Dinnon, K. H., 3rd; Leist, S. R.; Schafer, A.; Nakajima, N.; Takahashi, K.; Lee, R. E.; Mascenik, T. M.; Edwards, C. E.; Tse, L. V.; Boucher, R. C.; Randell, S. H.; Suzuki, T.; Gralinski, L. E.; Kawaoka, Y.; Baric, R. S., SARS-CoV-2 D614G Variant Exhibits Enhanced Replication ex vivo and Earlier Transmission in vivo. bioRxiv 2020.

22. Kathy Leung, Y. P., Gabriel M Leung, Tommy T. Y. Lam, Joseph T. Wu, Empirical transmission advantage of the D614G mutant strain of SARS-CoV-2. medRxiv 2020.09.22.20199810; doi: https://doi.org/10.1101/2020.09.22.20199810, 2020.

23. Volz, E.; Hill, V.; McCrone, J. T.; Price, A.; Jorgensen, D.; O’Toole, A.; Southgate, J.; Johnson, R.; Jackson, B.; Nascimento, F. F.; Rey, S. M.; Nicholls, S. M.; Colquhoun, R. M.; da Silva Filipe, A.; Shepherd, J.; Pascall, D. J.; Shah, R.; Jesudason, N.; Li, K.; Jarrett, R.; Pacchiarini, N.; Bull, M.; Geidelberg, L.; Siveroni, I.; Consortium, C.-U.; Goodfellow, I.; Loman, N. J.; Pybus, O. G.; Robertson, D. L.; Thomson, E. C.; Rambaut, A.; Connor, T. R., Evaluating the Effects of SARS-CoV-2 Spike Mutation D614G on Transmissibility and Pathogenicity. Cell 2021, 184 (1), 64–75 e11.

24. Plante, J. A.; Liu, Y.; Liu, J.; Xia, H.; Johnson, B. A.; Lokugamage, K. G.; Zhang, X.; Muruato, A. E.; Zou, J.; Fontes-Garfias, C. R.; Mirchandani, D.; Scharton, D.; Bilello, J. P.; Ku, Z.; An, Z.; Kalveram, B.; Freiberg, A. N.; Menachery, V. D.; Xie, X.; Plante, K. S.; Weaver, S. C.; Shi, P. Y., Spike mutation D614G alters SARS-CoV-2 fitness. Nature 2020.

25. Matsuyama, S.; Nagata, N.; Shirato, K.; Kawase, M.; Takeda, M.; Taguchi, F., Efficient activation of the severe acute respiratory syndrome coronavirus spike protein by the transmembrane protease TMPRSS2. J Virol 2010, 84 (24), 12658–64.

26. Gobeil, S. M.; Janowska, K.; McDowell, S.; Mansouri, K.; Parks, R.; Manne, K.; Stalls, V.; Kopp, M. F.; Henderson, R.; Edwards, R. J.; Haynes, B. F.; Acharya, P., D614G Mutation Alters SARS-CoV-2 Spike Conformation and Enhances Protease Cleavage at the S1/S2 Junction. Cell Rep 2021, 34 (2), 108630.

27. Benton, D. J.; Wrobel, A. G.; Xu, P.; Roustan, C.; Martin, S. R.; Rosenthal, P. B.; Skehel, J. J.; Gamblin, S. J., Receptor binding and priming of the spike protein of SARS-CoV-2 for membrane fusion. Nature 2020, 588 (7837), 327–330.

28. Starr, T. N.; Greaney, A. J.; Hilton, S. K.; Ellis, D.; Crawford, K. H. D.; Dingens, A. S.; Navarro, M. J.; Bowen, J. E.; Tortorici, M. A.; Walls, A. C.; King, N. P.; Veesler, D.; Bloom, J. D., Deep Mutational Scanning of SARS-CoV-2 Receptor Binding Domain Reveals Constraints on Folding and ACE2 Binding. Cell 2020, 182 (5), 1295–1310 e20.

29. Pettersen, E. F.; Goddard, T. D.; Huang, C. C.; Couch, G. S.; Greenblatt, D. M.; Meng, E. C.; Ferrin, T. E., UCSF Chimera--a visualization system for exploratory research and analysis. J Comput Chem 2004, 25 (13), 1605–12.

30. Case, D. A.; Belfon, K.; Ben-Shalom, I. Y.; Brozell, S. R.; Cerutti, D. S.; III, T. E. C.; Cruzeiro, V. W. D.; Darden, T. A.; Duke, R. E.; Giambasu, G.; Gilson, M. K.; Gohlke, H.; Goetz, A. W.; Harris, R.; Izadi, S.; S.A. Izmailov, K. K., A. Kovalenko, R. Krasny, T. Kurtzman, T.S. Lee, S. LeGrand, P. Li, C. Lin, J. Liu, T. Luchko, R. Luo, V. Man, K.M. Merz, Y. Miao, O. Mikhailovskii, G. Monard, H. Nguyen, A. Onufriev, F. Pan, S. Pantano, R. Qi, D.R. Roe, A. Roitberg, C. Sagui, S. Schott-Verdugo, J. Shen, C.L. Simmerling, N.R. Skrynnikov, J. Smith, J. Swails, R.C. Walker, J. Wang, L. Wilson, R.M. Wolf, X. Wu, Y. Xiong, Y. Xue, D.M. York and P.A. Kollman AMBER 2020. University of California, San Francisco 2020.

31. Maier, J. A.; Martinez, C.; Kasavajhala, K.; Wickstrom, L.; Hauser, K. E.; Simmerling, C., ff14SB: Improving the Accuracy of Protein Side Chain and Backbone Parameters from ff99SB. J Chem Theory Comput 2015, 11 (8), 3696–713.

32. Jorgensen, W. L.; Chandrasekhar, J.; Madura, J. D.; Impey, R. W.; Klein, M. L., Comparison of Simple Potential Functions for Simulating Liquid Water. J Chem Phys 1983, 79 (2), 926–935.

33. Socher, E.; Sticht, H.; Horn, A. H. C., The conformational stability of nonfibrillar amyloid-beta peptide oligomers critically depends on the C-terminal peptide length. ACS Chem Neurosci 2014, 5 (3), 161–7.

34. Salomon-Ferrer, R.; Götz, A. W.; Poole, D.; Le Grand, S.; Walker, R. C., Routine Microsecond Molecular Dynamics Simulations with AMBER on GPUs. 2. Explicit Solvent Particle Mesh Ewald. J Chem Theory Comput 2013, 9 (9), 3878–88.

35. Götz, A. W.; Williamson, M. J.; Xu, D.; Poole, D.; Le Grand, S.; Walker, R. C., Routine Microsecond Molecular Dynamics Simulations with AMBER on GPUs. 1. Generalized Born. J Chem Theory Comput 2012, 8 (5), 1542–1555.

36. Le Grand, S.; Götz, A. W.; Walker, R. C., SPFP: Speed without compromise-A mixed precision model for GPU accelerated molecular dynamics simulations. Comput Phys Commun 2013, 184 (2), 374–380.

37. Berendsen, H. J. C.; Postma, J. P. M.; Vangunsteren, W. F.; Dinola, A.; Haak, J. R., Molecular-Dynamics with Coupling to an External Bath. J Chem Phys 1984, 81 (8), 3684–3690.

38. Ryckaert, J. P.; Ciccotti, G.; Berendsen, H. J. C., Numerical-Integration of Cartesian Equations of Motion of a System with Constraints - Molecular-Dynamics of N-Alkanes. J Comput Phys 1977, 23 (3), 327–341.

39. Hopkins, C. W.; Le Grand, S.; Walker, R. C.; Roitberg, A. E., Long-Time-Step Molecular Dynamics through Hydrogen Mass Repartitioning. J Chem Theory Comput 2015, 11 (4), 1864–74.

40. Roe, D. R.; Cheatham, T. E., 3rd, PTRAJ and CPPTRAJ: Software for Processing and Analysis of Molecular Dynamics Trajectory Data. J Chem Theory Comput 2013, 9 (7), 3084–95.

41. Söldner, C. A.; Horn, A. H. C.; Sticht, H., Interaction of Glycolipids with the Macrophage Surface Receptor Mincle - a Systematic Molecular Dynamics Study. Sci Rep 2018, 8 (1), 5374.

